# The ARF GAPs ELMOD1 and ELMOD3 act at the Golgi and Cilia to Regulate Ciliogenesis and Ciliary Protein Traffic

**DOI:** 10.1101/2021.09.15.460558

**Authors:** Rachel E. Turn, Yihan Hu, Skylar I. Dewees, Narra Devi, Michael P. East, Katherine R. Hardin, Tala Khatib, Joshua Linnert, Uwe Wolfrum, Michael J. Lim, James E. Casanova, Tamara Caspary, Richard A. Kahn

## Abstract

ELMODs are a family of three mammalian paralogs that display GTPase activating protein (GAP) activity towards a uniquely broad array of ADP-ribosylation factor (ARF) family GTPases that includes ARF-like (ARL) proteins. ELMODs are ubiquitously expressed in mammalian tissues, highly conserved across eukaryotes, and ancient in origin, being present in the last eukaryotic common ancestor. We described functions of ELMOD2 in immortalized mouse embryonic fibroblasts (MEFs) in the regulation of cell division, microtubules, ciliogenesis, and mitochondrial fusion. Here, using similar strategies with the paralogs ELMOD1 and ELMOD3, we identify novel functions and locations of these cell regulators and compare them to those of ELMOD2, allowing determination of functional redundancy among the family members. We found strong similarities in phenotypes resulting from deletion of either *Elmod1* or *Elmod3* and marked differences from those arising in *Elmod2* deletion lines. Deletion of either *Elmod1* or *Elmod3* results in the decreased ability of cells to form primary cilia, loss of a subset of proteins from cilia, and accumulation of some ciliary proteins at the Golgi, predicted to result from compromised traffic from the Golgi to cilia. These phenotypes are reversed upon expression of activating mutants of either ARL3 or ARL16, linking their roles to ELMOD1/3 actions. Thus, we believe that ELMOD1 and ELMOD3 perform multiple functions in cells, most prominently linked to ciliary biology and Golgi-ciliary traffic, and likely acting from more than one cellular location.

## INTRODUCTION

Eukaryotic cells rely upon cell signaling, or the integration of intracellular and extracellular cues, both to ensure their survival and to thrive. A well-known class of cell signaling proteins includes the regulatory GTPases. Despite identifying these regulators through their conserved GTPase domains, we lack detailed models for how each paralog works in cells to mediate discrete functions. One central, technical challenge is the fact that specificity of GTPase actions is often lost in in vitro assays using purified components, as they fail to replicate the intricate cellular contexts under which they drive biological functions. Other challenges include that GTPases frequently act from multiple sites yet localize to each site only transiently. This holds true for GTPases along with the guanine nucleotide exchange factors (GEFs^1^) that activate them and the GTPase activating proteins (GAPs) that both inactivate the GTPase and frequently serve effector functions. The ability of proteins to recruit transiently to discrete cellular locations to drive cellular functions is clearly exemplified by the ARF GAPs (1–5). Most ARF GAPs can perform dual functions due to their recruitment to specific sites by the activated GTPases to which they bind, and in turn recruit other proteins/effectors. At least some of these recruited factors can regulate GTP hydrolytic (GAP) activity, thus terminating the upstream signal while propagating the downstream signal. Together, this system provides both spatial and temporal resolution to finely hone signal output. This functional duality for ARF GAPs led to an increased appreciation of the need to understand their roles in cells (1,3).

The ARF family of regulatory GTPases is large, with 30 mammalian members that are highly conserved throughout eukaryotic evolution, 16 of which are predicted to have been present in the last eukaryotic common ancestor (6). ARFs and ARLs regulate a range of essential cellular processes including roles in membrane traffic, primary cilia, mitochondria, tubulin biogenesis, and microtubule dynamics (1,7–10). Unfortunately, we have only fragmentary data on the substrate (GTPase) specificity for any ARF GAP (e.g., in vitro GAP activity using purified proteins) or ARF GEF as well as the cellular functions of each (1,3,10–14). There are 28 known or predicted ARF GAPs, 24 of which were identified based on the presence of an ARF GAP domain (15–17). Of the ARF GAPs that lack the consensus ARF GAP domain, three mammalian proteins share an ELMO domain (ELMOD1-3) with broad specificity for GTPases in the ARF family (18), while the sole outlier, RP2, possesses GAP activity specific to ARL2 and ARL3 (19). Those ARF GAPs that share the ARF GAP domain have been tested using in vitro assays and found to act with varying degrees of specificity towards one or more of the six mammalian ARFs (ARF1-6; though typically only ARF1 and ARF6 have been tested), but few have been tested against any of the 22 ARF-like (ARL) GTPases. In efforts to gain insights into signaling by the ARLs, we purified ELMOD2 as an ARL2 GAP and found that all three of the human ELMOD proteins act on several different ARLs as well as ARFs (18,20). Thus, this broad specificity in in vitro GAP assays suggests the possibility that ELMODs act in an even broader set of signaling pathways than do the ARF GAPs.

The ELMO (cell EnguLfment and MOtility) family of proteins consists of six members in humans that share an ELMO domain and are equally divided into structurally and phylogenetically distinct subgroups, termed ELMOs and ELMODs (18,21–24). The ELMODs are ancient proteins that span the entire diversity of eukaryotes and date back to the last eukaryotic common ancestor (18). In contrast, the ELMOs are a more recent evolutionary family, found only in metazoans and fungi, and are predicted to have emerged from the ELMODs (18). While the ELMODs are GAPs with broad substrate specificities towards ARFs and ARLs (18,20,25,26), the ELMOs primarily function in complex with DOCK proteins as unconventional GEFs for RHO/RAC GTPases (which act at the leading edge of migrating cells (21,27–30)) and lack any GAP activity. In MEFs, ELMOD2 serves critical roles in the regulation of cytokinesis, ciliation, microtubule stability, mitochondrial fusion, ciliogenesis, anchoring of rootlets to centrosomes, and lipid metabolism at lipid droplets (25,31–35). Effects of ELMOD2 deletion on mitochondrial fusion and microtubules are linked to ARL2 (25,34), while effects on cell division/abscission are specifically suppressed by expression of activated ARF6 (25), demonstrating that one ELMOD can regulate multiple pathways through distinct GTPases. Because *Elmod2* deletion in MEFs results in so many phenotypes, it is challenging to identify root causes of each and the extent of connectedness among them. Furthermore, because ELMOD2 is a member of a three gene family in mammals, sharing a common functional domain, we sought to compare these actions to those of other family members. This is both to pave the way for defining the cellular functions of each of the family members and also to assess the extent of functional redundancy that may suppress the appearance of even more deficits in cells when any one member is mutated or deleted.

While the three mammalian ELMODs share broad substrate specificities for GTPases in the ARF family, they display different specific activities for those substrates (20), giving them great potential to play important clinical roles if mutated. For example, mutations in *ELMOD2* are linked to familial idiopathic pulmonary fibrosis as well as antiviral response in both mice and humans (36,37). In contrast, *ELMOD1* mutations and *ELMOD3* mutations are both linked to deafness, autism, and intellectual disability in humans and mice (38–45). Hearing deficits linked to mutation of either *Elmod1* or *Elmod3* in mice are also linked to defects in stereocilia, or actin-based apical projections of inner ear hair cells. For stereocilia to form, the hair cell must first project a primary cilium (in the context of hair cells, called the “kinocilium”) which plays a critical role in signaling for proper stereocilia morphology and orientation. Almost all mammalian cells generate at least one cilium, motile (*e.g.*, flagellum) or non-motile (primary cilia and kinocilia), and some cells (particularly epithelial) display multiciliation (46–49). Cilia are primarily present in cells that have exited the cell cycle at least in part because centrosomes are involved in formation of the basal body, the associated distal appendages, the ciliary pocket, and the microtubule-based axoneme that extends and pushes out the nascent cilium. The early steps of ciliogenesis, termed licensing, can be induced in cell culture via serum starvation. This initiates the recruitment of CEP164 to the mother centriole followed by recruitment of TTBK2 (50–53) and later “uncapping” or loss of CP110 (54–57). The release of CP110 is followed by ciliary vesicle docking and axoneme extension (57). For additional details of ciliogenesis, consult the following reviews (58–60). Once formed, cilia are dependent upon three known protein complexes to engineer the entry and export of ciliary proteins as well as movement along the axoneme: intraflagellar transport complex A (IFT-A, retrograde traffic), IFT complex B (IFT-B, anterograde traffic), and the BBSome (61–71). Despite characterizations of their actions as multi-subunit protein complexes, there is evidence that one of the 16 components of the IFT-B complex, IFT20, acts independently of the complex at the Golgi to promote traffic to the ciliary base (72,73). Knockout (KO) of IFT-20 causes a more severe phenotype than does KO of the IFT-A core subunit IFT140 (74).

Much less is known about routes proteins take to get to or through the ciliary base, but there are likely several different routes and regulators as well as differences between membrane and soluble proteins. Soluble proteins <~100 kDa (9 nm diameter) can simply diffuse into cilia (75–77). On the other hand, membrane proteins are synthesized on the rough ER membrane, traffic through the Golgi, and are directed to the basal body through incompletely understood processes. Notable among the proteins implicated in regulating import and/or retention of ciliary proteins are four ARF family GTPases: ARL3, ARL6, ARL13B, and ARL16 (7). ARL6 acts in concert with the BBSome (78). Together, they regulate export of G protein coupled receptors from cilia (61,79,80). ARL13B interacts with INPP5E, a PIP 5'-phosphatase, and PDE6D (aka PDE6δ or Pr/BPδ), a transporter of prenylated cargos that includes INPP5E (81–83). ARL13B possesses GEF activity for ARL3, which in turn can bind directly to PDE6D resulting in the release of its cargo (84–87). While ARL13B can be palmitoylated on cysteine residues near the N-terminus, and thereby become anchored in the membrane, it is not clear how ARL3 is retained in cilia, though its local activation by ARL13B may regulate its localization in cilia. We also have recently identified defects in ciliogenesis and ciliary protein content in cells deleted for ARL16 (S.I.D and R.A.K., manuscript in preparation). Even less well understood are the regulators of these ciliary GTPases, as to date very little is known about the ARF GEFs and GAPs that work in primary cilia.

Here we describe the use of a number of approaches, relying heavily on CRISPR/Cas9 edited MEFs, to begin defining the cellular functions of *Elmod1* and *Elmod3* and comparing the degree of functional overlap/specificity among the family members (25,34,35). ELMOD1 and ELMOD3 studies to date have focused on in vitro biochemical activities (18,20,26) and analyses of mutations in mammals (38–45,88), so the cellular functions remain largely uncharacterized. We use MEFs as our model system to allow detailed comparisons to earlier studies of *Elmod2* KO in isogenic cell lines. We include descriptions of organelle morphologies and pathways that appear unaltered in these KO lines as a broad scan for unexpected, potential functions, as such data also provide supportive evidence of the specificity of the defects identified.

## RESULTS

### *CRISPR/Cas9 knockouts of* Elmod1, Elmod3, *or both in immortalized mouse embryonic fibroblasts*

In efforts to identify the cellular functions of ELMOD1 and ELMOD3, we used CRISPR/Cas9 in MEFs to introduce frameshift mutations in *Elmod1*, *Elmod3*, or both *Elmod1/Elmod3*. We previously used the same procedure to generate *Elmod2* KO MEFs and described multiple resulting phenotypes (25,34,89). We used two or more guides to target exons (see Fig. S1A, B and Methods for details) and identified at least two clones from each guide to obtain lines with null alleles. We summarize the alleles for all cell lines in Fig. S1C. We predict that all 17 lines result in the loss of functional proteins or null alleles, as these frameshifts are targeted upstream of the sole functional (ELMOD) domain. Thus, we refer to these lines as knockouts (KOs) and double KO (*Elmod1/Elmod3* DKO) lines. We also generated clones with no mutations in the targeted region. These are retained as additional controls, as they went through the same transfection, selection, and cloning process as the KO and DKO lines. We refer to such lines as CRISPR WT (CWT) to distinguish from the parental WT lines. The use of more than one guide, multiple alleles, and independent clones helps alleviate concerns regarding off-target effects, alternative splicing, and downstream initiation (89,90). As an additional protection against off-target effects, we also performed rescue experiments in which we transiently re-expressed the disrupted gene to test for reversal of key phenotypes.

### *Many organelles and processes appear unaltered in cells deleted for* Elmod1, Elmod3, *or* DKOs

Because ELMODs display in vitro GAP activity with a number of different ARF family GTPase substrates, known to regulate a wide array of cellular processes, we screened the KO lines using markers of different organelles and processes to identify gross changes. For example, we previously reported (18) localization of ELMOD1 at nuclear speckles, but no evidence of changes in nuclear speckles were evident in any of these KO lines (Fig. S2A). Hoechst staining revealed no evidence of changes in nuclear size, number, or morphology (Fig. S2B). Because we first purified ELMODs as ARL2 GAPs, we looked at processes previously reported to be regulated by ARL2 but found no evident changes in mitochondrial morphology (Fig. S2C), microtubule networks (Fig. S2D), or centrosome numbers (25,35). Phalloidin staining for actin was not overtly altered among *Elmod1* or *Elmod3* KO cell lines (Fig. S2E) when compared to WT cells. Because we earlier identified Golgi as a site of action of over-expressed ELMOD1 in HeLa cells (18), we looked at markers of the cis-Golgi (GM130; Fig. S2F), TGN (Golgin-97; Fig. S2G), TGN/endosomes (BIG2/ARFGEF2; Fig. S2G), and recycling endosomes (RAB11-FIP1, RAB11-FIP3, or RAB11-FIP5), and again found no evidence of gross morphological alterations. Because KO of *Elmod2* caused a number of readily identified phenotypes, we also examined those in *Elmod1/3* KO cells. Cell cycle was not obviously altered based upon propidium iodide staining and DNA content analysis using flow cytometry (Fig. S3A). Neither were ciliary rootlets altered, as they were intact and appeared similar in size around centrosomes that were not significantly more separated than centrosomes in WT cells (Fig. S3B, C), in contrast to what was observed in *Elmod2* KOs (35). Cold and nocodazole sensitivity of microtubules are obvious in MEFs lacking ELMOD2 (25), but neither phenotype was evident in lines deleted for *Elmod1*, *Elmod3*, or both. Number and size of focal adhesions also are unaltered from WT in *Elmod1/3* KO and DKO cells (Fig. S4).

### ELMOD1 and ELMOD3 are localized in ciliary compartments of mouse photoreceptor cells

Previous data from our lab revealed novel functions for ELMOD2 as a negative regulator of ciliogenesis, as its deletion caused increases in ciliation. Motivated by this and the genetic data linking both ELMOD1 and ELMOD3 to defects in stereocilia, we investigated potential roles for ELMOD1 and ELMOD3 in cilia. To begin to test this model, we searched for novel localizations for ELMOD1 and ELMOD3. ELMODs are low abundance proteins, with ELMOD1 estimated to be present <0.01% of total cell protein in HeLa cells (18). Despite evidence of the expression of all three transcripts in MEFs from RNA-seq data (though levels of the mRNAs were so low as to challenge accurate quantification, C_t_ ≥35), they are each absent from a database of over 8,400 proteins identified in MEFs by mass spectrometry (unpublished observation), presumably due to low protein expression. Neither commercial nor in-house ELMOD1 and ELMOD3 antibodies yielded specific signal by immunofluorescence in MEFs. We next turned to the well-studied model for proteins implicated in cilia, retinal photoreceptor cells, to identify specific localization of each protein.

Mouse retinas were prepared and processed as described previously (35,91,92), and in Materials and Methods. In cryosections through the retina of eGFP-Centrin mouse, eGFP-Centrin was visualized to identify the connecting cilium and basal body (93). Counterstaining with DAPI was used to demonstrate the nuclear layers. ELMOD1 (Fig. 1A) and ELMOD3 (Fig. 1B) were found in the plexiform layers of the retina (Fig. 1, IPL), where a high density of synapses localize, though each of the ELMODs are more prominent in the photoreceptor region (Fig. 1; IS, CC, OS). Higher magnification of this region revealed that both ELMOD1 and ELMOD3 were present at the base of the connecting cilium (Fig. 1A, B, lower panels) which links the outer segments (= modified primary cilium) and inner segments of photoreceptor cells and resembles an elongated transition zone of primary cilia (94). There, ELMOD1 staining was found in the basal body and the daughter centriole, as well as in between the basal body and the daughter centriole (Fig. 1A, lower panel, C). In contrast, we found ELMOD3 not only in the linkage between the basal body and the daughter centriole but also in an extension of the connecting cilium towards the axoneme of the photoreceptor outer segment (Fig. 1B, lower panel, D), which is consistent with the previously reported localization of ELMOD2 (35). There, in the basal part of the outer segment, the *de novo* formation of the photo-sensitive disc membranes of the outer segment occurs. This process is highly regulated by cilia-specific molecules and is powered by actin polymerization (95–97).

**Figure 1:**
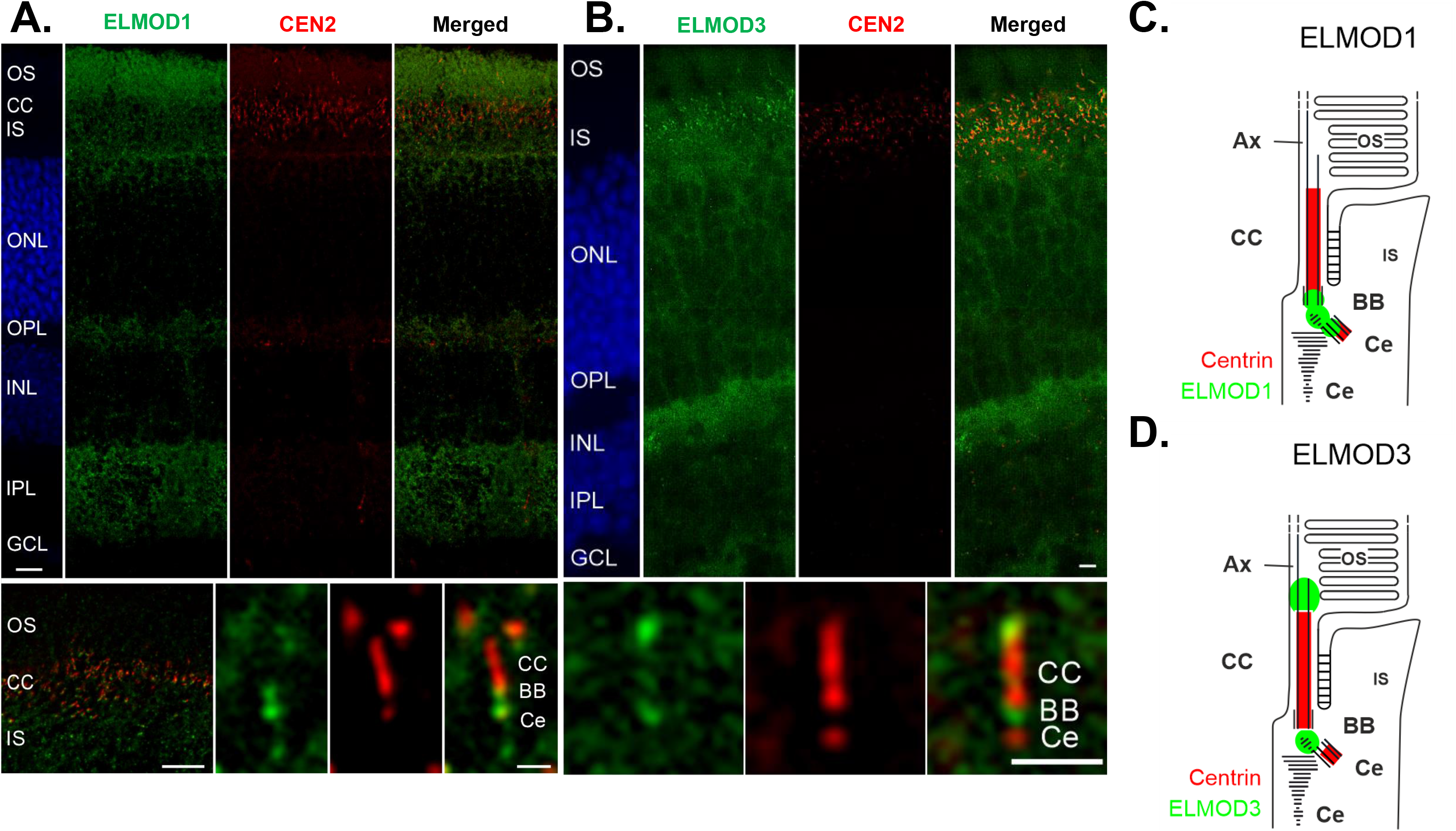
ELMOD1 and ELMOD3 show overlapping, but also some discrete, localization patterns in mouse retinal photoreceptor cells. Retinae of eGFP-CETN2 mice were processed and analyzed with a deconvolution microscope, as described under Materials and Methods, to determine ELMOD1 and ELMOD3 localization. (**A,** large upper panel) ELMOD1 immunolabeling of cryosections through the retina revealed a prominent staining in a region of the photoreceptor cells comprising the outer segment (OS), the connecting cilium (CC, red, Cen 2), and the inner segment (IS). In addition, fade staining was observed in the outer and inner plexiform layer (OPL, IPL) while other retinal layers, the outer and inner nuclear layer (ONL, INL, blue DAPI staining), and ganglion cell layer (GCL) did not show substantial staining. (lower panel) Higher magnification of photoreceptor region (left) and of the ciliary part of a photoreceptor cell (right) revealed ELMOD1 localization at the base of the connecting cilium (CC) in the basal body (BB) and the adjacent daughter centriole (Ce) (both red) as well as within the bridge in between. (**B,** large upper panel) ELMOD3 immunolabeling in a transgenic eGFP-CETN2 mouse retina revealed staining at the region containing photoreceptor cells and both plexiform layers (OPL, IPL) (lower panel). Higher magnification of the ciliary part of a photoreceptor cell showed that ELMOD3 was restricted to a localization between the BB and Ce at the base of the CC. In addition, ELMOD3 was stained at the base of the outer segment, a compartment above the CC. (**C, D**) Scheme of ELMOD1 and ELMOD3 photoreceptor cells. Scale = A, B, upper panels: 15 μm; lower panels (higher magnifications), 5 μm.

In conclusion, ELMOD1 and ELMOD3 show distinct localization patterns but overlap in their localization at the base of the connecting cilium. Both the base as well as the tip of the connecting cilium are associated with numerous signaling pathways. Interestingly ELMOD1, ELMOD2, and ELMOD3 all show joint localization around the connecting cilium. This could indicate a common site of action for the three proteins at the base, and for ELMOD2 and ELMOD3 at the tip of the connecting cilium.

### Both Elmod1 *KOs and* Elmod3 *KOs cause decreased ciliogenesis at a late step in licensing*

Given the evidence for ELMOD1 and ELMOD3 localization to basal bodies in retinal cells and roles of ELMOD2 in cilia, we next examined whether primary cilia form at normal rates and with typical protein content in *Elmod1* and *Elmod3* KO lines. Using acetylated tubulin (Ac Tub) antibody to mark the axonemes, we monitored the percentage of ciliated cells after 24, 48, and 72 hours of serum starvation (0.5% FBS). We found ciliation was strongly <u>decreased</u> in *Elmod1* and *Elmod3* single KO MEFs, with <10% of cells on average having a cilium after 24 hrs of serum starvation, compared to >60% in WT controls (Fig. 2A, B). While the percentage of ciliated cells increased at later times of serum starvation in both WT populations and each of the KO clones, the latter never approached levels seen in WT cells (Fig. 2A). In the course of this work, we visually inspected hundreds of cells in biological replicates from multiple clones for each genotype and observed no obvious, consistent differences in ciliary length.

**Figure 2:**
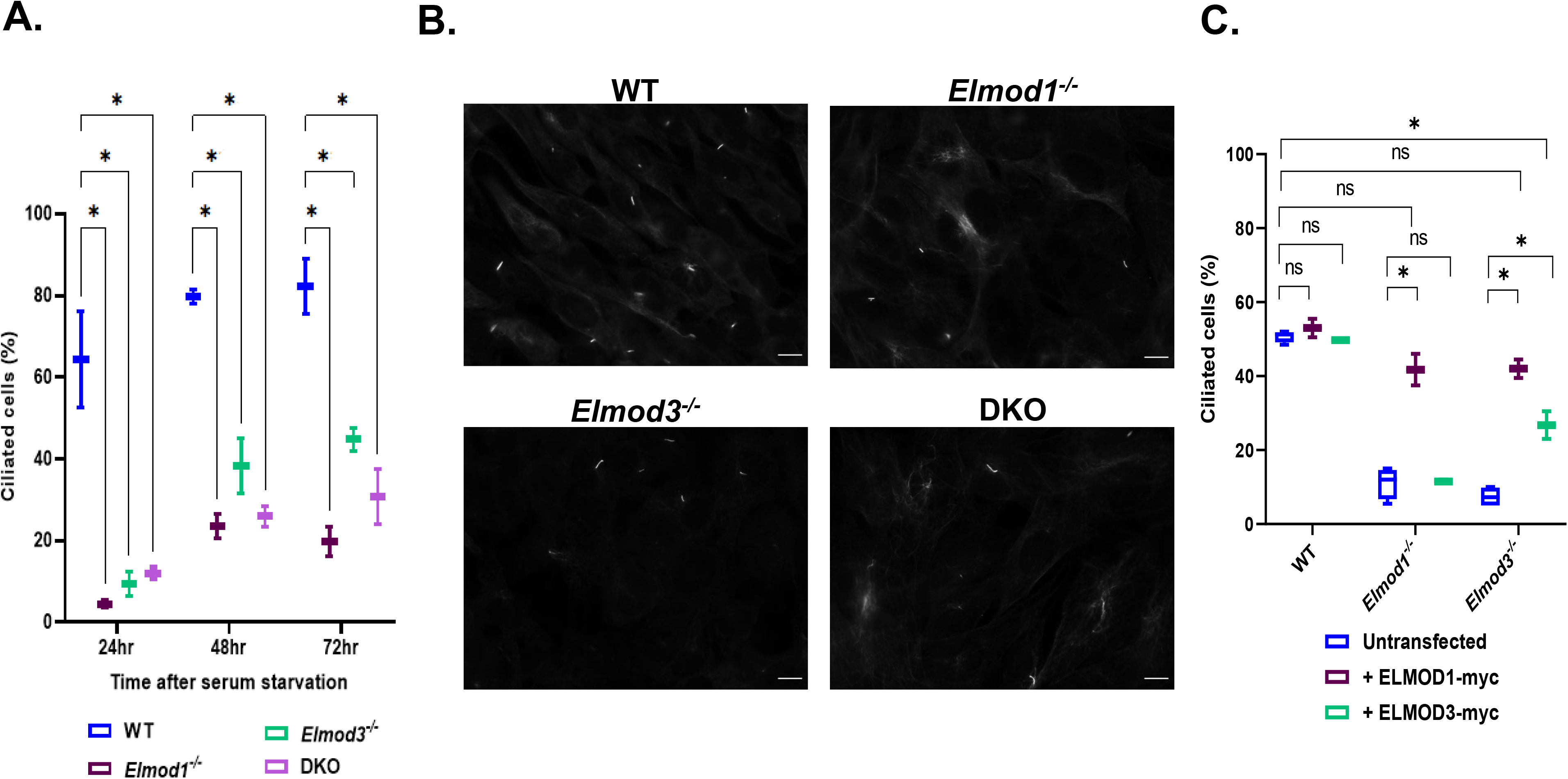
*Elmod1* and/or *Elmod3* KOs in MEFs cause decreased ciliation. Cloned lines of WT, *Elmod1* KO, *Elmod3* KO, and DKO MEFs were serum starved for either 24, 48, or 72 hours to induce ciliogenesis prior to staining for Ac Tub (to mark cilia) as described under Materials and Methods. **(A)** The percentage of cells with cilia were scored (2 cell lines per genotype, 100 cells each) at each time point. The experiment was performed in duplicate, and all values were averaged, with error bars indicating the SEM. **(B)** Representative images of WT and KO cells were collected at 60x magnification by widefield imaging, 72 hours after serum starvation. Scale bar = 10 μm. **(C)** Plasmids directing expression of ELMOD1-myc, ELMOD3-myc, or GFP were transiently transfected into WT, *Elmod1* KO or *Elmod3* KO lines (two each). One day later, they were serum starved for 24 hrs and then stained and scored for cilia as in panel **A**. Only cells with evident protein expression (myc positive) were scored. Experiments were scored in duplicate, 100 cells per replicate. Results were plotted using GraphPad. Statistical significance was assessed via One-Way ANOVA. ns = not significant, * = p<0.05.

We performed rescue experiments in which we transiently expressed either ELMOD1-myc or ELMOD3-myc proteins in KO, DKO, and WT MEFs and scored ciliation after 24 hrs of serum starvation. Expression of ELMOD1-myc in *Elmod1* KO lines was sufficient to bring ciliation percentages close to those of parental WT controls, while expression of ELMOD1-myc in WT cells had no significant effect on ciliation (Fig. 2C). Because we scored all myc positive cells, even those staining weakly, there may be a lower limit of expression required to bring ciliation percentages fully up to those seen in WT cells. Expression of ELMOD3-myc was also tested and found capable of reversing the decreased ciliation in *Elmod3* deleted cells, though only about halfway back to levels seen in WT cells (Fig. 2C). Interestingly, expression of ELMOD1-myc also rescued ciliation in cells deleted for *Elmod3* (Fig. 2C). Indeed, the effect of ELMOD1-myc expression on restoration of ciliation in *Elmod3* KO lines was comparable to that seen in ELMOD1-myc rescue of *Elmod1* KOs and of ELMOD3-myc in *Elmod3* KO cells (Fig. 2C). In contrast, expression of ELMOD3-myc in *Elmod1* KOs failed to restore ciliation (Fig. 2C). We compared the relative levels of ELMOD1-myc and ELMOD3-myc expression by immunoblotting for the myc tag in total cell lysates of WT cells 24 hr after transient expression of each protein (Fig. S5). This revealed that ELMOD1-myc is expressed to considerably higher levels than is ELMOD3-myc (Fig. S5), perhaps explaining its greater potency in rescue. Unfortunately, the low levels of endogenous ELMOD protein expression in MEFs prohibited quantitative comparisons between endogenous and exogenous protein expression.

Having identified that loss of ELMOD1 and ELMOD3 leads to ciliation defects, we next sought to monitor the early steps required for licensing of ciliogenesis in mutant MEFs. We previously showed that ELMOD2 regulates the earliest stages of ciliogenesis, so we predicted that ELMOD1 and/or ELMOD3 may also be acting early in ciliogenesis. We examined recruitment of CEP164 (normally to nascent distal appendages) and loss of CP110 or “uncapping,” both of which are critical for licensing of axoneme growth (50,54,56,57,98,99). We found no differences in the degree of either CEP164 recruitment (Fig. S6A) or CP110 loss (Fig. S6B) in any of the mutant MEF lines. We also stained cells for CEP290, a marker of the transition zone at the base of cilia, and also found no changes in the extent of its recruitment (Fig. S6C). Together, we interpret these results as evidence that the ciliation defect seen in *Elmod1/3* KO lines is downstream of the uncapping of CP110 at the distal appendages and of transition zone construction involving CEP290.

### *KO of* Elmod1 *or* Elmod3 *alters the protein composition in cilia*

We next sought to determine if loss of ELMOD1 or ELMOD3 also had effects on ciliary protein composition. To monitor the protein content of cilia, we first examined ARL13B, which is bound to the ciliary membrane through N-terminal palmitoylation and localizes along the entire ciliary length (7,100–103). We observed reduced ciliary ARL13B staining in both KO and the DKO lines, revealing compromised import (or increased export) in cells that lack ELMOD1 and/or ELMOD3 (Fig. 3A,B). In WT cells, all Ac Tub staining overlapped with ARL13B staining, indicating that WT cilia contain ARL13B, with little variation in the intensity of staining. In contrast, each of the *Elmod1*, *Elmod3,* and DKO lines examined displayed a large majority of Ac Tub positive cilia in which staining of ARL13B was clearly reduced and was even undetectable in ~15-20% of cilia (Fig. 3A). These differences were so stark, at least in part due to the strong staining of ciliary ARL13B in WT cells, that the quantification of pixel intensities was unnecessary. In DKO cells, more cilia (~26%) were devoid of ARL13B, though the difference in levels from the single KO lines was not statistically significant.

**Figure 3:**
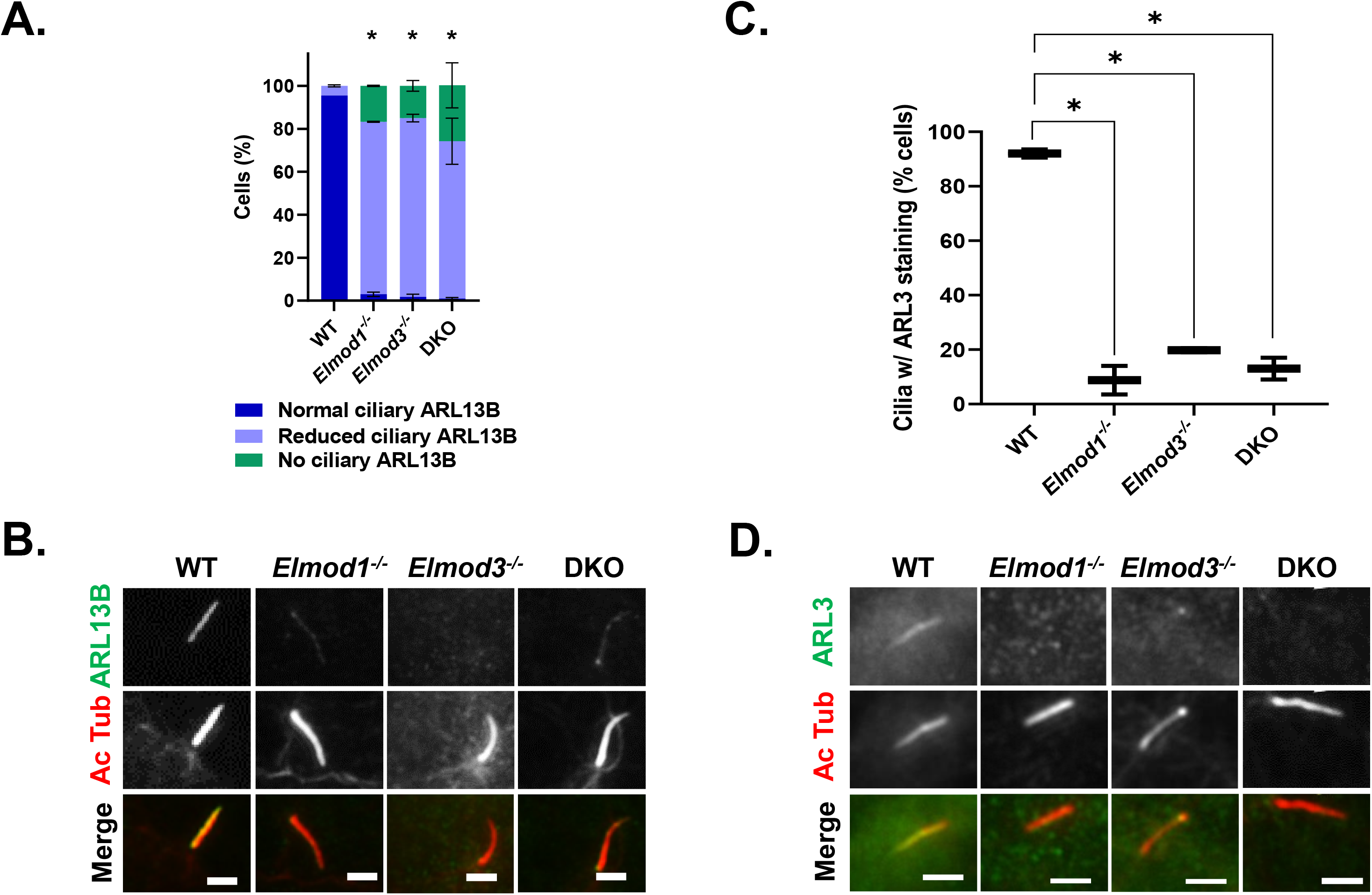
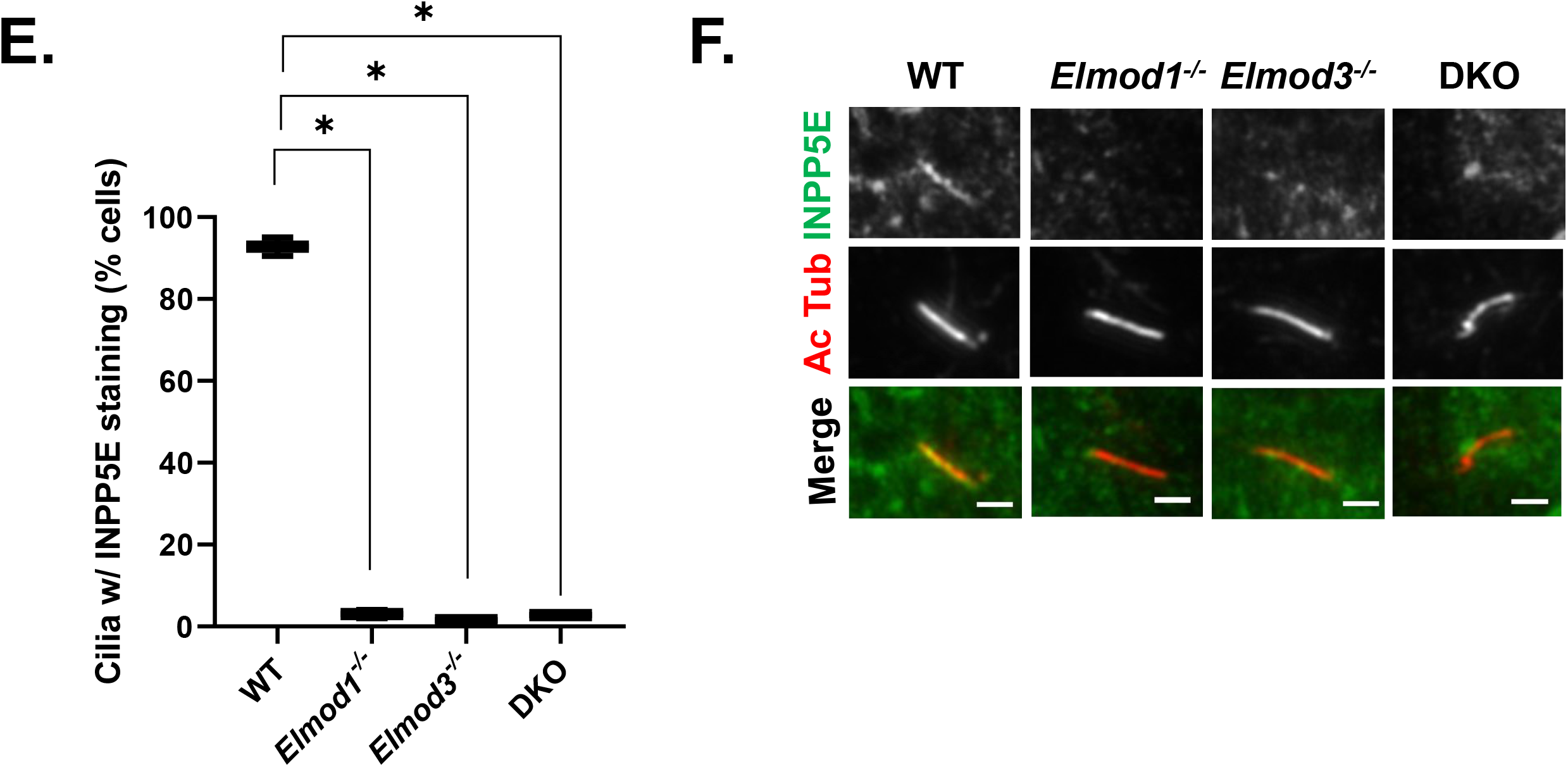
*Elmod1* or *Elmod3* KO causes loss of ARL13B, ARL3, and INPP5E from cilia. Cells (two lines per genotype, from different guides) were grown to ~80% confluence before inducing ciliation via serum starvation for 72 hrs, and then staining for Ac Tub and either ARL3, ARL13B, or INPP5E. **(A)** ARL13B levels were binned as normal (readily identifiable prior to confirming with the Ac Tub channel), reduced (only identifiable as ciliary upon confirmation with Ac Tub staining), or absent (no staining evident that overlaps with Ac Tub). Scoring was performed on 100 cells for each of two cell lines in replicate, and error bars represent SEM. (**B**) Cells treated as in (A), stained for Ac Tub and ARL13B, were collected by widefield microscopy at 100X, and representative images are shown. Note that in these images the *Elmod1* KO and DKO cells were scored as reduced, as faint staining is evident, while in the *Elmod3* KO cell it is not. **(C)** Ciliary ARL3 was scored in cells treated as described in panel A. Due to the weaker overall staining of ARL3, scoring was binned as either present or absent from cilia, identified via Ac Tub staining. Scoring was performed on 100 cells for each of two cell lines in replicate, and box-and-whisker plots from the scoring are shown; error bars = SEM. (**D**) Representative images from cells stained for ARL3 and Ac Tub are shown. **(E)** INPP5E and Ac Tub staining was performed as described under Materials and Methods, and scoring of INPP5E presence in cilia was performed as described in panel C, being binned as either present or absent. (**F**) Representative images from cells stained for Ac Tub and INPP5E are shown. Statistical significance for all data was determined via One-Way ANOVA using GraphPad Prism Software. * = p<0.05. For all images shown, scale bar = 10 μm.

ARL13B can act as a GEF for ARL3 (85–87). Import of ARL3 into cilia is thought to be mediated by simple diffusion, while its retention may be ARL13B-dependent (104,105). We examined ARL3 localization in cilia and found it to be strongly decreased in *Elmod1* and *Elmod3* KO lines (Fig. 3C,D). Because staining of ARL3 in cilia is not as intense as that of ARL13B, we scored in a binary fashion, “normal” versus “reduced,” with the latter being scored only when unambiguously below levels seen in most WT cells. Despite this conservative approach to scoring, the loss of ARL3 staining was quite strong, with over 80% of cilia having reduced ARL3 staining, compared to WT cells where fewer than 10% displayed lower ciliary ARL3. Thus, both ARL13B and ARL3 are each strongly reduced in cilia, either due to decreased import or increased export (failed retention).

Among other actions in cells, ciliary ARL3 binds the prenylated cargo transporter PDE6D, causing it to release its cargo at that site (86,106,107). Perhaps the best known prenylated cargo involved is the lipid phosphatase INPP5E, active in modifying the lipid composition of cilia (104,108–114). Once in cilia, INPP5E is thought to bind directly to ARL13B, which aids in its retention (81–83). Thus, we also analyzed ciliary content of INPP5E in *Elmod1*, *Elmod3*, and DKO cells. We found a near complete loss of INPP5E staining in each of the KO lines (Fig. 3E, F). The reduction in INPP5E in cilia may be an indirect result from the loss of ARL13B or ARL3, or it may result from other defects (see below).

Cilia are required to transduce vertebrate Hedgehog (Hh) signaling, which is regulated by both ARL13B and INPP5E (110,115–118). GLI3 is one of three transcription factors that localize to cilia and increase at the ciliary tip in response to Hh stimulation (119). We examined ciliary GLI3 in the KO lines and found no differences in GLI3’s ciliary staining or enrichment at the ciliary tip (Fig. S7). We also stained for markers of the intraflagellar transport complexes IFT-A (IFT140) and IFT-B (IFT88) and found no differences between WT and KOs in their staining in cilia (Fig.S8). Taken together, these data show that ciliary protein content is altered in cells lacking *Elmod1* and/or *Elmod3,* but it appears to be a select group of proteins with previously identified ciliary links that are reduced in cilia.

### INPP5E and IFT140 accumulate in the Golgi in Elmod1/3 and DKO MEFs

As a C-terminal isoprenylated protein, INPP5E traffics throughout the cell as cargo of the PDE6D transporter and is deposited onto membranes upon either ARL2 or ARL3 binding to PDE6D (86,104,106,113,120–122). Thus, the absence of INPP5E from cilia might be explained by the lack of ARL3, if required for dislocation from PDE6D, or lack of retention offered by binding to ARL13B (81,82). While staining ELMOD1/3 KO lines for INPP5E, we noticed an increase in INPP5E staining intensity in serum starved cells (24 hrs) that co-localized with the Golgi marker GM130 (Fig. 4A,B). Under these conditions, INPP5E staining at Golgi in WT cells was weak and usually absent. In contrast, the presence of INPP5E at Golgi is evident in well over half of all *Elmod1*, *Elmod3*, and DKO cell lines after 24 hrs of serum starvation (Fig. 4A,B). Note that INPP5E staining at cilia was performed using PFA as fixative, while its presence at Golgi was only evident after methanol fixation. Interestingly, by 72 hrs this effect was reduced, with only ~25-30% of *Elmod1*, *Elmod3*, or DKO cell displaying INPP5E at Golgi. This reduction over time suggests a transience to this phenomenon that may result from a pulse of new INPP5E synthesis in response to serum starvation or a redistribution within the cell. A more detailed time course was not investigated at this time but may be an interesting future direction to pursue.

**Figure 4:**
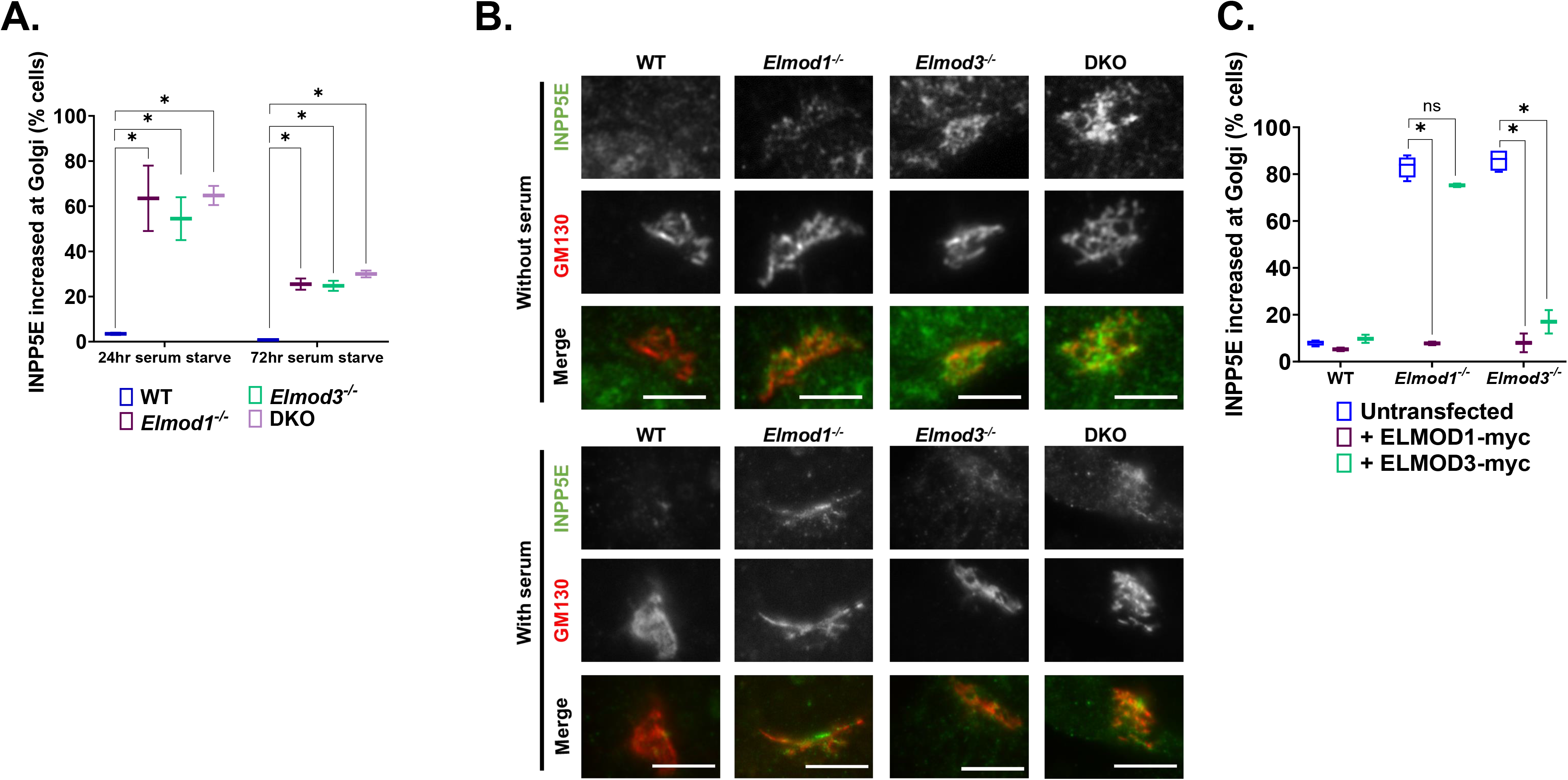
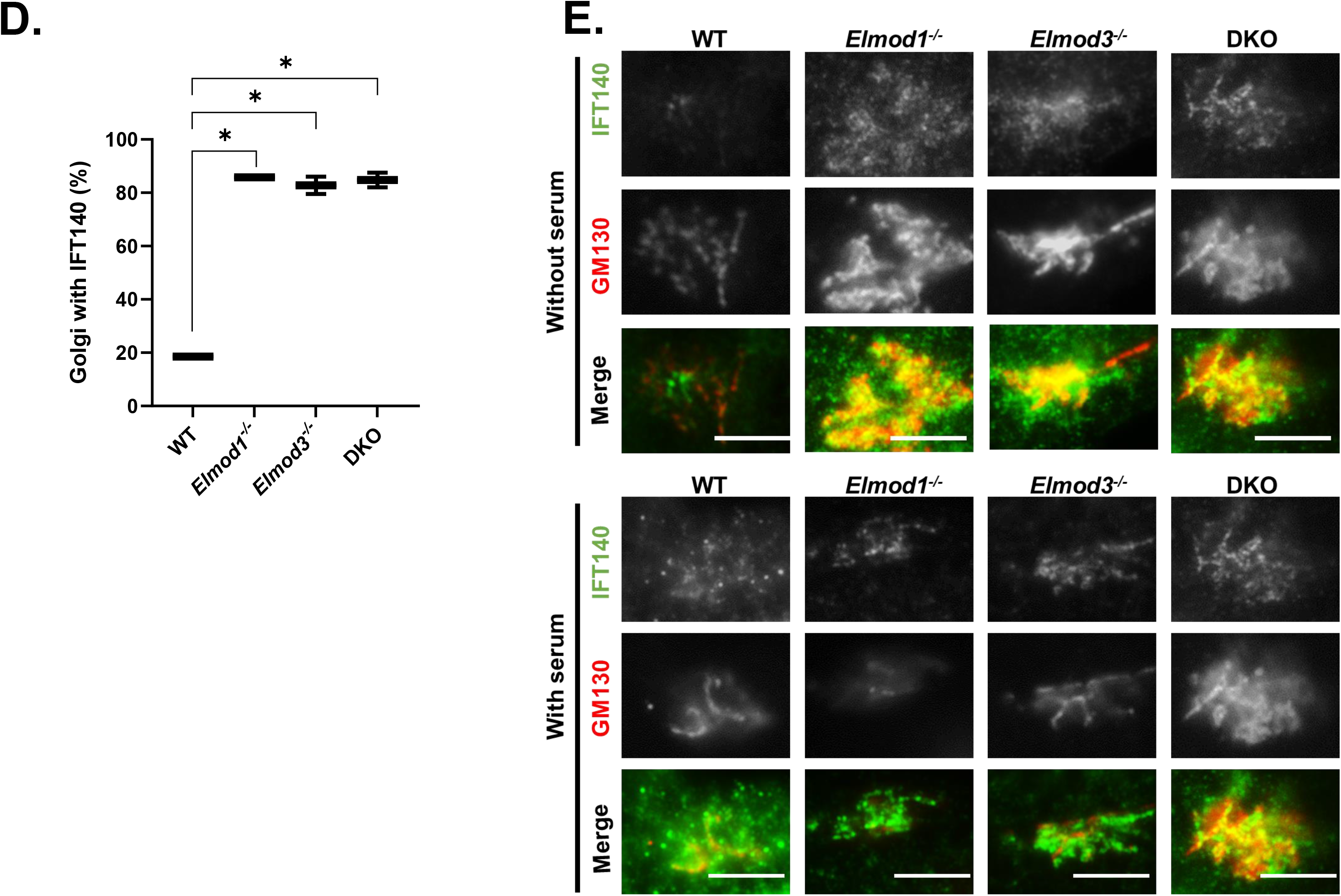
INPP5E and IFT140 accumulate at the Golgi in *Elmod1* or *Elmod3* KO lines. Cells were serum starved for 24 or 72 hrs, followed by staining for GM130 and either INPP5E (**A-C**) or IFT140 (**D-F**). (**A**) The presence of INPP5E co-localizing with GM130 was scored for each genotype, using two different clones of each and scoring 100 cells in duplicate experiments. Error bars = SEM. * = p<0.05. (**B**) Representative images of WT and KO cells serum-starved for 72 hr and stained for GM130 and INPP5E were collected at 100x magnification with widefield microscopy. Scale bar = 10 μm. **(C)** WT, *Elmod1*, and *Elmod3* KO cells were transfected with plasmids directing the expression of ELMOD1-myc or ELMOD3-myc. 24 hrs later, the cells were serum starved for 24 hr before fixing and staining for GM130 and INPP5E. The increased abundance of IFT140 at Golgi was scored as in panel A. Error bars = SEM. * = p<0.05. (**D**) The presence of IFT140 co-localizing with GM130 was scored for each genotype, using two clones of each knock out line and one WT line, scoring 100 cells in duplicate experiments. Error bars = SEM. * = p<0.05. (**E**) Images of IFT140 at Golgi, using GM130 as marker, in each line. Cells were serum starved for 24h and stained for GM130 and IFT140.

We performed rescue experiments using transient expression of either ELMOD1-myc or ELMOD3-myc. Expression of either protein had no impact on the percentages of cells displaying staining of INPP5E at the Golgi in WT cells (Fig. 4C). Expression of either protein in lines deleted for that protein reduced the staining of INPP5E at the Golgi to near WT levels (Fig. 4C). As we observed for rescue of overall ciliation percentages, expression of ELMOD1-myc reversed the increased levels of INPP5E both in *Elmod1* and *Elmod3* KO lines, while ELMOD3-myc expression only rescued this phenotype in *Elmod3* KOs (Fig. 4C).

The increased localization of INPP5E at Golgi in *Elmod1* and *Elmod3* KO lines also prompted us to look for the presence of other ciliary proteins at Golgi. We found that IFT140, a core component of IFT-A, is also increased at the Golgi in *Elmod1/3* KO lines compared to WT cells (Fig. 4D,E). In marked contrast, we did not detect any increased staining for other IFT proteins at the Golgi, including IFT81, IFT88, or IFT144 (data not shown). Thus, while the presence of IFT140 at the Golgi is increased in KO lines, it appears to be there independently of the IFT-A complex. The increased Golgi staining of IFT140 was also evident in KO lines without serum starvation (Fig. 4E), suggesting a block or delay in traffic that is independent of induced ciliogenesis.

In the process, we also observed strong IFT140 staining that was typically adjacent to, but rarely overlapped with, the Golgi marker (GM130) and was more restricted in space. Co-staining of IFT140 and rootletin in WT cells demonstrates that IFT140 is present at ciliary rootlets and that this likely is the source of the strong staining near the Golgi (Fig. 4F). IFT140 staining of rootlets was unaltered from that seen in WT cells in any of the *Elmod1/3* KO lines (Fig. S9).

### *Expression of either activated ARL3 or activated ARL16 rescues ciliation defects and accumulation of INPP5E and IFT140 at Golgi in* Elmod1 *KO*, Elmod3 *KO*, and DKO *lines*

ARF GAPs have dual functions in cells: both to terminate/dampen signaling from specific GTPases to which they bind and to propagate the downstream signal, most commonly via recruitment of other proteins (1,3). In efforts to identify the GTPase(s) acting in a GAP-sensitive pathway, it has proven fruitful to test for rescue of phenotypes resulting from deletion of the GAP using activated mutants of different GTPases. Because the commonly used “Q to L” mutants (corresponding to Q71 in ARF1 or Q61 in HRAS) are often quite toxic in cells, we use “fast cycling” mutants that become activated independently of an ARF GEF and, unlike the Q to L mutants, can still cycle between active and inactive conformations and thereby propagate the signal without locking up the pathway (25,35,123). Transient transfections were used to express fast cycling mutants of ARF1 (ARF1[T161A]-HA), ARF5 (ARF5[T161A]-HA), ARL3 (ARL3[L161A]-myc), or ARL16 (ARL16[R153A]-myc) in WT and KO MEFs. Expression of either activated ARL3-myc or ARL16-myc restored ciliation percentages near WT levels (Fig. 5A), functionally linking ARL3 and ARL16 to the actions of ELMOD1 and ELMOD3 in ciliation. In contrast, neither ARF1[T161A]-HA nor ARF5[T161A]-HA expression had any effect on ciliation of WT, *Elmod1,* or *Elmod3* KO cells (Fig. 5A), supporting the conclusion that there is specificity to the GTPases acting with ELMOD1/3 in ciliation. Finally, we examined whether the expression of the same two activated ARF family members, ARL3 and ARL16, that restored ciliation also reversed the accumulation of INPP5E at the Golgi. We found that each of these GTPases partially reversed the increased staining of INPP5E at the Golgi (Fig. 5B). Thus, deficiencies in ciliation correlate well with increased INPP5E localization at Golgi, and each is reversed either by expression of deleted ELMOD or by activated ARL3 or ARL16, suggesting functional linkage between these two lesions.

**Figure 5:**
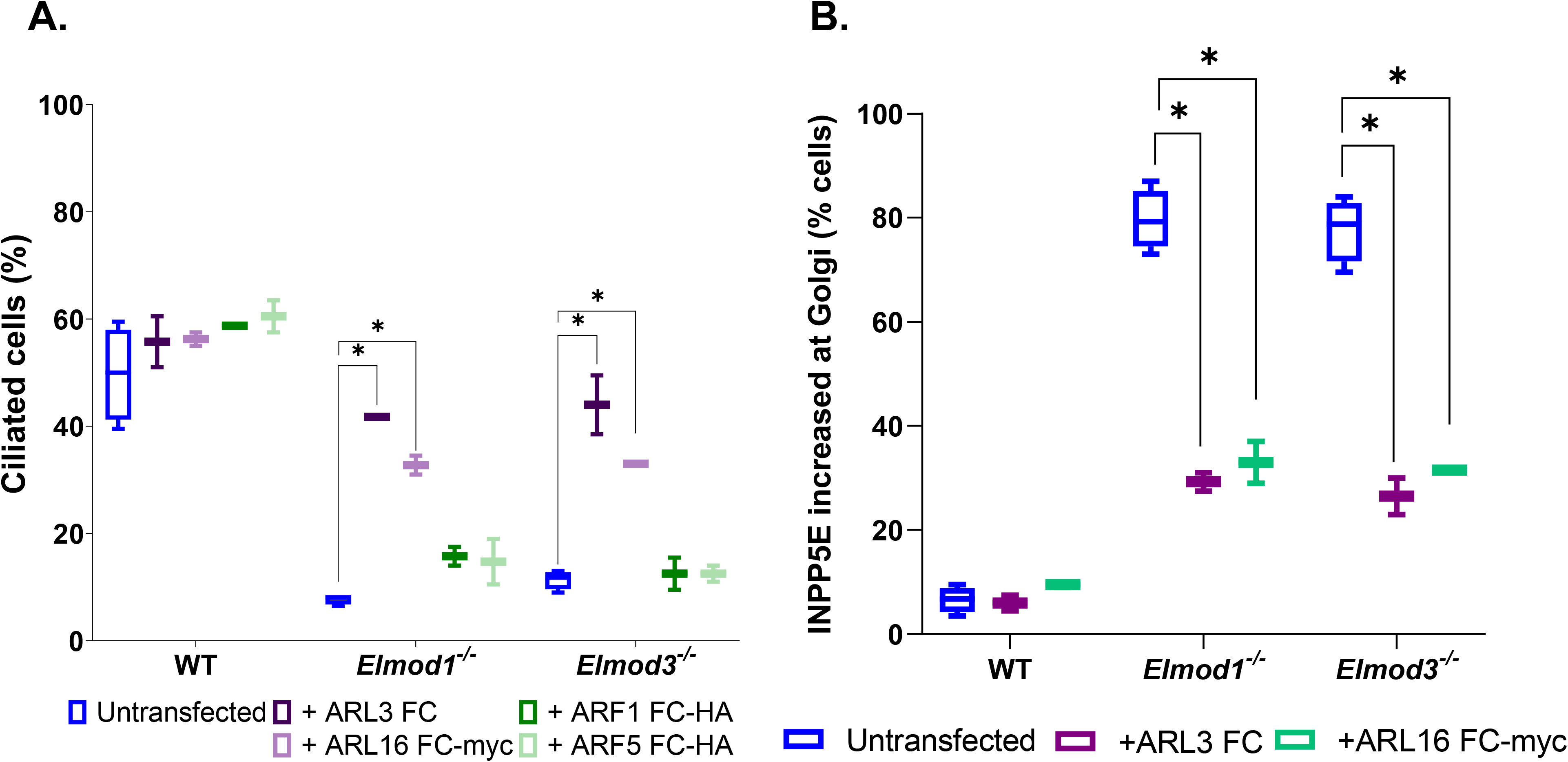
The ciliation defect in *Elmod1* or *Elmod3* KO cells can be reversed upon transient expression of activated ARL3 or ARL16. (**A**) WT, *Elmod1* KO, and *Elmod3* KO lines were transfected with plasmids directing expression of fast cycling mutants of ARF1-HA, ARF5-HA, ARL3, or ARL16-myc. The next day, cells were serum starved for 24 hours and then fixed and stained for Ac Tub and the corresponding tag. (**B**) The same procedure was carried out as described for panel A, except that only fast cycling mutants of ARL3 and ARL16 were examined and that staining was for the expressed protein and INPP5E. Only those cells expressing exogenous proteins were scored. Two lines for each genotype were scored, 100 cells each, and the experiment was repeated in duplicate. The box-and-whisker plot shows the averages, with error bars representing the SEM. * = p <0.05.

## DISCUSSION

We developed KO MEF lines to define cellular roles of ARF GAPs ELMOD1 and ELMOD3 in mammalian cells. We found that both *Elmod1* and *Elmod3* KOs have decreased ciliation, loss of import or retention of several ciliary proteins that are important in signaling, and accumulation of some of the same proteins at Golgi. We also identified likely roles for ARL16 and ARL3 in ELMOD1 and ELMOD3 function. In several respects, these results parallel studies of the other paralog, ELMOD2, in that ELMOD1 and ELMOD3 appear to act at more than one place and in concert with more than one GTPase within a given cell (25,34,35). However, ELMOD1 and ELMOD3 play distinct roles from those described for ELMOD2, most notably with *opposite* effects on ciliation, as summarized in Table I. We propose a model whereby ELMOD1 and ELMOD3 act at cilia, most likely the basal body, to regulate ciliogenesis and protein import, and also at the Golgi, to regulate the export of specific cargos (INPP5E and IFT140) that may strongly influence ciliary generation and protein content (Fig. 6). These findings clearly call for future studies to better define molecular mechanisms of all three ELMODs in ciliogenesis, ciliary protein recruitment, Golgi traffic, and related processes in multiple tissues and cell types.

**Table 1:** Summary of phenotypes found in MEFs deleted for *Elmod1* KO, *Elmod2* KO, *Elmod3* KO, and *Elmod1/Elmod3* DKO MEFs using CRISPR-Cas9. The top of the graph indicates the cell lines, the left most column lists the phenotypes assayed and markers used, and phenotypes are indicated under each cell line as normal (no change from WT) or other.

| **<u>Phenotype</u>** | **<u>*Elmod1* KO</u>** | **<u>*Elmod3* KO</u>** | **<u>DKO</u>** | **<u>*Elmod2* KO</u>** |
| --- | --- | --- | --- | --- |
| Mitochondrial morphology (HSP60 staining) | Normal | Normal | Normal | Fragmented |
| Centrosome number (γ-tubulin staining) | Normal | Normal | Normal | Supernumerary |
| DNA content (Propidium iodide staining) | Normal | Normal | Normal | Polyploid |
| Nuclear number (Hoechst staining) | Normal | Normal | Normal | Multinucleated |
| Rootlet morphology (Rootletin staining) | Normal | Normal | Normal | Fragmented |
| Microtubule sensitivity (Cold/nocodazole) | Normal | Normal | Normal | Increased |
| Ciliation (% of cells) (Ac Tub staining) | Decreased | Decreased | Decreased | Increased |
| Multiciliation) (Ac Tub staining) | Normal | Normal | Normal | Multiciliated |
| Ciliogenesis licensing (Cep164, CP110 staining) | Normal | Normal | Normal | Increased (CEP164↑<br>CP110↓) |
| ARL13B/ARL3 in cilia (ARL13B/ARL3 staining) | Decreased | Decreased | Decreased | Normal |
| INPP5E in cilia (INPP5E staining) | Decreased | Decreased | Decreased | ND |
| IFT140 at basal body (IFT140 staining) | Decreased | Decreased | Decreased | ND |
| IFT140 at Golgi (IFT140 staining) | Increased | Increased | Increased | ND |

**Figure 6:**
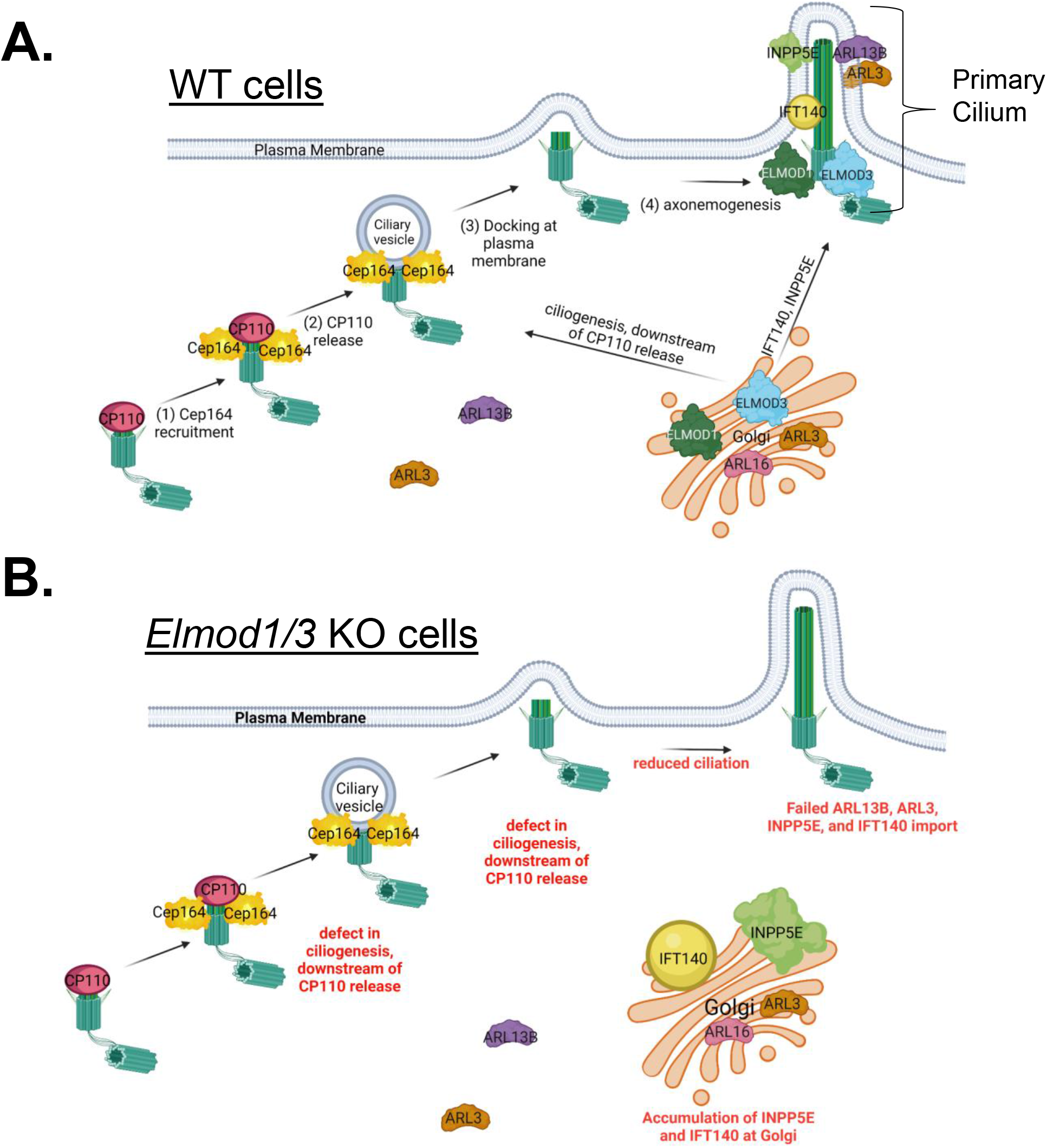
Model for ELMOD1 and ELMOD3 function as regulators of ciliogenesis and traffic of key ciliary cargoes. We propose that ELMOD1 and ELMOD3 are acting in concert and in at least three processes to regulate ciliogenesis and the traffic of key ciliary cargoes from the Golgi to cilia. With respect to ciliogenesis, we propose that ELMOD1 and ELMOD3 are regulating ciliogenesis from the basal body, after the release of CP110 from distal appendages. ELMOD1 and ELMOD3 are also required for ARL13B in cilia, which in turn aids in the ciliary retention of ARL3. These first two actions may be closely linked in space, though neither ARL13B nor ARL3 are required for ciliogenesis, so we consider them separate at this time. We believe that ELMOD1 and ELMOD3 also act from the Golgi to regulate export of INPP5E and IFT140, though through distinct mechanisms, because export of INPP5E requires PDE6D while export of IFT140 does not. Finally, we speculate that ELMOD1 and ELMOD3 can also act from endosomes, perhaps directly on ARL16, and that in their absence ARL16 is strongly increased on endosomal membranes. Perhaps this would result in its depletion from other sites, including Golgi, causing delays or defects in export of specific proteins from the Golgi. Figure created using BioRender.

The diversity in sites of action and effects on different processes makes dissection of molecular mechanisms for ELMOD1 and ELMOD3 a challenge-a common issue for members of the ARF family and their regulators, as recently highlighted (1). To begin to address such questions, we looked at localization of ELMOD1 and ELMOD3 in cultured mammalian cells and retinal tissue. We reported earlier that endogenous ELMOD1 is overwhelmingly soluble by cell fractionation of normal rat kidney (NRK) cells (18). Unfortunately, neither homemade nor commercial antibodies directed against ELMOD1 and ELMOD3 were sensitive enough to identify locations of endogenous proteins in MEFs. However, we found ELMOD1 at the basal body and centriole of mouse retinal cells (Fig. 1A,C). ELMOD3 staining overlapped with that of ELMOD1 at the base of the connecting cilium of the centriole in photoreceptor cells (Fig. 1B,D). The overlap in locations between ELMOD1 and ELMOD3 in mouse retinal cells is consistent with previously reported localizations of ELMOD1 in hair cells at the cell apex (40) and ELMOD3 in stereocilia and kinocilia in the organ of Corti and inner ear (39). The localizations of ELMOD3 are also quite similar to what we reported earlier for ELMOD2 in mouse retina (35). Previous work revealed that endogenous ELMOD3 localizes to stereocilia and kinocilia in both inner and outer hair cells (39) and that GFP-ELMOD3 recruits to stereocilia in inner ear explants and to cortical actin structures in a confluent monolayer of MDCK cells (41). ELMOD3 has been implicated as a target of the cilia-related RFX2 transcription factor (124) and was also localized to basal bodies in a related, high throughput screen (125). Thus, the locations of ELMOD1 and ELMOD3 in cells and tissues are incompletely understood and may show tissue specificity. We fully expect the ELMODs to act at additional sites, likely in all cells, *e.g.*, at subcortical actin, as suggested by expression of ELMOD3-GFP in MDCK cells (43). Based on our data, though, we predict that these players are commonly found at the basal body/transition zone where they may act on aspects of ciliogenesis and regulated protein import/export. Interestingly, we also note strong staining of ELMOD1/3 at the inner plexiform layer of the retina which is rich in synapses. These findings may lay the groundwork for future studies designed to look at potential roles for ELMODs at synapses in neural tissues.

We describe three cilia-related consequences from loss of either ELMOD1 or ELMOD3 that likely represent distinct actions of these ARF GAPs: decreased ciliation, loss of several proteins from cilia, and increased accumulation of some of the same proteins at the Golgi. We found that the defect in ciliation occurs at a step downstream of “uncapping” of the mother centrosome. Potential later steps where they might act include the expansion of the ciliary pocket or extension of the axoneme, particularly as ARF family GTPases play multiple roles in vesicular membrane traffic (126–131), and ARL2 specifically in regulation of microtubules (132–139). In contrast, the other paralog ELMOD2 is required quite early in the process, upstream of CEP164 recruitment, and its absence results in <u>increases</u> both in the percentage of cells that ciliate but also in multiciliation, which were not phenotypes resulting from KO of either ELMOD1 or ELMOD3. Furthermore, ELMOD2 localizes to rootlets, and its deletion causes striking fragmentation of ciliary rootlets with centrosome separation (35). In contrast, neither *Elmod1* nor *Elmod3* KOs show such effects on rootlet integrity and centrosome separation (Fig. S3B,C). Together, these data point to a combination of discrete and overlapping functions between members of the ELMOD family.

We also discovered changes in the ciliary content of at least three proteins in cells lacking ELMOD1 and/or ELMOD3: ARL13B, ARL3, and INPP5E. Note that this is not a universal defect in all ciliary protein traffic, as such factors as Ac Tub, IFT88, IFT140, and GLI3 show normal localization, as do the regulators of early ciliogenesis (*i.e.*, CEP164 and CP110). The proteins that are lost are not all transported to cilia by a common mechanism, so a single site of action seems unlikely. ARL3, at ~20 kDa, is small enough to diffuse across the transition zone (75,104,140), and its abundance in cilia may be increased as a result of its binding/activation by ARL13B with its ARL3 GEF activity (86). If this is the case, then the loss of ARL3 may be simply secondary to the decreased ciliary abundance of ARL13B. INPP5E is a farnesylated cargo that is trafficked to cilia by transporter PDE6D, which releases cargos upon binding to activated ARL3 (104). However, ciliary targeting of INPP5E has been shown to be dependent upon ARL13B and not ARL2 or ARL3 (82,83). Thus, the decreased ciliary accumulation of INPP5E may also be an indirect result from the loss of ARL13B. How ARL13B gets to and is retained at cilia is less clear, though a role for Ahi1 acting at the transition zone to affect ARL13B levels has been shown, as have links to Tulp3 and RPGRIP1L (141–143). ARL13B becomes membrane-associated after palmitoylation of cysteines near its N-terminus and likely acts more like a membrane protein than the others (100,102,103). While mutations in ARL13B can cause decreased ciliation and shortened cilia, none of the other proteins with altered localization in *Elmod1/3* KO lines are required for growth of a cilium of normal length (144). Thus, we posit that the <u>traffic</u> defect(s) in *Elmod1* and *Elmod3* KOs, whether at the Golgi or ciliary base, represent a distinct lesion from that which decreases ciliogenesis overall, while the decrease in percentage of ciliated cells after serum starvation in *Elmod1/3* KO lines may be common to the losses observed in *Arl13b* null cells (101,102,117).

INPP5E co-localizes extensively with β-COP at the Golgi in 3T3-L1 adipocytes (145) and Tera-1 cells (146). In contrast, in RPE-hTERT cells, INPP5E was found at the ciliary axoneme with only minimal non-ciliary staining (112). INPP5E is cytosolic and at the plasma membrane in macrophages (147). Thus, while farnesylated INPP5E is thought to traffic throughout the cell while bound to PDE6D, just when and where these two first encounter one another is unknown. The accumulation of INPP5E and IFT140 at the Golgi in *Elmod1* and *Elmod3* KO cells may have physiologically important consequences to IFT-A assembly and functions as well as the regulation of phosphatidylinositol phosphates on multiple membranes, making clean dissection of molecular mechanisms challenging, but certainly worthy of further study. That is, the increased abundance of INPP5E at the Golgi and its loss from cilia are expected to cause alterations in the lipid composition of membranes at each compartment, with likely indirect consequences to localization and actions of any number of other proteins.

The finding that IFT140 and INPP5E accumulate at the Golgi in both *Elmod1* and *Elmod3* KO cells could explain their loss from cilia. We can detect INPP5E staining at the Golgi in WT MEFs, though it is clearly much fainter and less consistently present at Golgi than in *Elmod1/3* KO lines. This is consistent with a model in which IFT140 normally traffics through the Golgi, despite the lack of an identifiable membrane binding motif. Little is known about how IFT140 is moved through cells and assembles into the core IFT-A complex. In contrast to INPP5E, IFT140 has not previously been shown to localize to Golgi. Future studies into how IFT140 is trafficked will provide critical new insight into the mechanisms by which IFT-A import occurs-an understudied question.

These findings should raise awareness of the potential for proteins acting at the Golgi to have consequences in ciliary biology. A clear precedent has been set by studies of IFT20, which localizes to the basal body as well as the Golgi, where it works with GMAP210 in the sorting and traffic of proteins to ciliary membranes (72,148). The actions of IFT20 and IFT140 were compared and while each was found to be critical for membrane protein traffic to cilia, IFT20 was found to act at the Golgi while IFT140 acted from the plasma membrane (74). In this case, though, it was likely the IFT-A complex and not IFT140 acting alone that was under study, as IFT140 deletion would remove both pools. Interestingly, IFT20 has more recently been found to play a role in integrin recycling, cell migration, and focal adhesion dynamics, potentially linking these membrane dynamics to ciliary protein traffic in ways that call for further exploration (149). Open questions include how IFT140 associates with membranes (as it lacks a transmembrane domain as well as classic acylation and prenylation motifs), as well as how and where it assembles into the IFT-A core complex.

ELMOD1 and ELMOD3 display differences in specific activities against ARFs and ARLs in the in vitro GAP assay (20), yet almost identical phenotypes when KO’d in MEFs. The simplest model to emerge from these results is that they act in a common pathway, though perhaps on different steps. This is also supported by the findings that deletion of *Elmod1* <u>and</u> *Elmod3* in the same cells does not result in stronger phenotypes than either KO alone. That expression of ELMOD1-myc rescues the loss of ciliation in both *Elmod1* and *Elmod3* KO lines is also consistent with them sharing a common pathway and possibly with ELMOD1 acting downstream of ELMOD3, though the fact that ELMOD1-myc is expressed to higher levels than ELMOD3-myc may provide an alternate explanation for this finding. The partial rescue of *Elmod3* KO lines by ELMOD3-myc, and perhaps even its inability to reverse the ciliation defect in *Elmod1* KO lines, may result from lower levels of expression or interference in activity by the C-terminal tag, though the latter appears unlikely given the success of ELMOD1-myc in rescue. The finding that ELMOD3-myc expression incompletely rescues ciliation in *Elmod3* KOs and fails to rescue *Elmod1* KO line phenotypes may be due to lower levels of ELMOD3-myc expression compared to ELMOD1-myc. Both constructs contain codon-optimized open reading frames, and both use a common expression vector and the same epitope tag. Finally, we cannot exclude the possibility that while both proteins act on a common step or in a common pathway, they may have different specific activities as GAPs and thus require different levels of expression for full rescue.

The finding that *Elmod1* KO and *Elmod3* KO ciliary defects are each rescued by expression of either activated ARL3 or ARL16 is also consistent with ELMOD1/3 sharing a common pathway and strongly implicates these two GTPases in that pathway. The differences in functions identified for ELMOD1 and ELMOD3 vs ELMOD2 (Table I) likely result from their functional interactions with different ARF family GTPases: with ELMOD2 acting with ARL2 and ARF6 (25,34,35) and ELMOD1/3 acting with ARL3 and ARL16. A broader sampling of the 30 mammalian ARF family GTPases would be required to gain a more complete picture of the specificities for each of these GAPs/effectors in each pathway. ELMODs also have highly divergent N- and C-termini that are not thought to be involved in GTPase binding or hydrolysis. Yet, they may confer distinct spatial regulation or binding partners in cells. The recently released AlphaFold protein structure database (150,151) contains structure predictions for each of the three ELMODs from multiple species. While every ELMOD1 and ELMOD2 structure includes long and short α-helices at N- and C-termini, respectively, every ELMOD3 structure is predicted to have long (30-50 residues) unfolded regions that may lower protein stability as a result of greater access to proteases or binding to other proteins (that are *not* increased in expression). Despite such predicted structural or stability differences, we interpret rescue of phenotypes resulting from KO of an ARF GAP by an activating mutant of an ARF family GTPase as evidence of their interaction in a common pathway. Work from Cardenas-Rodriguez, et al (152) showed that, in zebrafish, mutations in *Cep290* can result in increased expression of a number of proteins, including ARL3, ARL13B, and UNC119 and that exogenous expression of these same proteins can reverse ciliary phenotypes. Unlike in this study in fish, we have not yet searched for changes in gene expression in response to KO of *Elmod*s in MEFs.

Our findings can also be considered in light of the evolutionary history of the ELMOD family. ELMODs are ancient proteins, with between one and six (in plants) genes present in a broad spectrum of eukaryotes; mammals express three, ELMOD1-3. While among the ELMODs some overlap in biochemical activities and shared specificities for GTPases exist (20), there are already some striking differences in the actions of ELMOD2 from those of ELMOD1 and ELMOD3 in MEFs, and likely all cells. The lack of changes in centrosome numbers and separation, rootlet integrity, cell cycle, and microtubule sensitivities in *Elmod1/3* KO cells is in marked contrast to what we recently reported in *Elmod2* KO lines (25,89) (Table I). The absence of changes in both CEP164 recruitment and release of CP110 is also opposite to those changes observed in ELMOD2 KO lines, which display large increases in the percentages of cells with >1 centrosome that stain positive for CEP164 and TTBK2 and negative for CP110 (Table I). Thus, ELMOD1/3 vs ELMOD2 each play roles in ciliogenesis but appear to act at distinct steps in the pathway, and their actions lead to opposite effects on ciliogenesis. Interestingly, a recent study in *Arabidopsis* identified roles for ELMODs in pollen aperture formation, with two paralogs apparently working together and another appearing to act in opposition to those two (153), perhaps analogous to what we report here. From our earlier phylogenetic analyses of the ELMODs, we concluded there was at least one, and likely two, ELMOD present in the last eukaryotic common ancestor (18). We speculate that the differences in functions ascribed to ELMOD2 vs ELMOD1/3 may represent a very early divergence in function within the ELMOD family in the evolution of eukaryotes.

Our characterization of ELMOD1 and/or ELMOD3 in mammalian cells has revealed a number of novel functions for these two ARF GAPs and also prompts new questions worth exploring. We focused on the use of MEFs, in large part to allow direct comparisons to comparable studies on ELMOD2, but recognize the possibility of differences between cell and tissue types. While we found that ciliation was compromised in *Elmod1/3* KO lines downstream of CP110 cap release, it is quite possible that detailed study, *e.g.*, using electron microscopy, may reveal defects in ciliary vesicle recruitment, axoneme extension, or other specific steps in ciliogenesis. Such studies are technically challenging and time consuming due to the small size of cilia, incomplete ciliation of most cell types, and need for serial sectioning. The finding that the increased abundance of INPP5E at Golgi was higher after 24 hr of serum starvation than after 72 hr is suggestive of a transience to this phenotype, though it was evident in both regular (10%) and low (0.5%) serum conditions. One possible explanation for this may be that export of INPP5E and IFT140 from the Golgi is normally occurring at very close to maximal rates and that loss of either ELMOD1 or ELMOD3 impairs that process, resulting in their accumulation. Finally, testing a more complete set of fast cycling mutants of the 30 mammalian ARF family GTPases for rescue of the phenotypes identified here would allow for a far better understanding of the specificities for GTPases by these GAPs in the pathways under study. We are currently working to generate such a complete collection of activating mutations, but this too will require more time and study.

## MATERIALS AND METHODS

### Reagents, antibodies, plasmids

All chemicals used were purchased from commercial sources. The following commercial antibodies were used in these studies: SC35 (1:500; Abcam; ab11826), HSP60 (1:1000; Stressgen; ADI-SPA-807), α-tubulin (1:1000; Sigma; T9026), γ-tubulin (1:5000; Sigma T6557 or Abcam ab11317), GM-130 (1:000; BD/Transduction 610823), Golgin97 (1:500; Proteintech; 12640-1-ap), BIG2 (1:200; EMD Millipore; MABS1246), Rootletin (1:500; EMD Millipore; ABN1686), centrin clone 20H5 (1:1000; Sigma; 04-1624), Ac Tub (1:1000; Sigma; T5192), α-myc (1:1000; Invitrogen R950-25 or Abcam ab9132), CEP164 (1:100; Santa Cruz; sc-515403), CP110 (1:100; Proteintech; 66448-1-ig), CEP290 (1:100; Proteintech; 22490-1-ap), ARL13B (1:1000; Proteintech; 17711-1-AP), INPP5E (1:100; Proteintech; 17797-1-ap), Gli3 (1:1000; R&D Systems; AF3690), β-COP (1:2,000; ThermoFisher PA1-061), IFT140 (1:500; Proteintech; 17460-1-AP) IFT88 (1:500; Proteintech; 13967-1-ap), α-HA (1:1000; Covance; MMS-101P), β1-integrin 1 (1:200; Sigma MAB1997), active integrin clone 9EG7 (1:200, BD-Pharmingen 553715). The following antibodies were generously provided from other labs: polyclonal, rabbit antibodies raised against FIP1 (1:500) and FIP5 (1:500) were from Rytis Prekeris (University of Colorado), and sheep anti-FIP3 (1:500) was from Jim Goldenring (Vanderbilt University). Our lab generated ARL3 rabbit polyclonal antibody, which we use at 1:1000 dilution (154).

Plasmids directing expression of mouse ELMOD1-myc or ELMOD3-myc were codon optimized for expression and synthesized by GeneArt, and later moved into the pcDNA3.1 vector. Fast cycling mutants of GTPases were generated by site directed mutagenesis of the residue corresponding to T161 in ARF6 (123), followed by sequencing of the complete open reading frame to confirm the mutation and lack of extraneous changes.

### Cell culture, transfections, and induction of ciliation

MEFs used in this study were originally obtained from the American Tissue Type Collection (ATCC CRL-2991) and were maintained in DMEM with 10% fetal bovine (FBS) serum (Atlanta Biologicals, S11150) and 2mM glutamine. Antibiotics were not used in routine culture to minimize chances of mycoplasma contamination, which was monitored by DNA staining. In any experiment in which comparisons were planned between lines with different genotypes, attention was paid to ensure comparable feeding/plating schedules and cell densities were used to minimize the likelihood that such variables may confound the data or their interpretations.

Transient transfections of WT or KO MEFs for rescue experiments were performed using jetOPTIMUS (VWR, 76299-634). Cells seeded at 90% confluence were transfected with a ratio of 4ug DNA: 4uL jetOPTIMUS transfection reagent: 400uL jetOPTIMUS buffer adhering to the protocol provided by the manufacturer. The DNA/jetOPTIMUS mixture was added dropwise to each respective well, and samples were returned to 37°C to incubate for 24 hours. Cells were maintained in normal growth serum (i.e., 10% FBS in DMEM). The next day, cells were replated as needed for different experiments.

Induction of ciliation involved switching to low FBS (0.5%) medium one day after plating and allowing ciliation to progress for 24-72 hours before fixing cells. Ciliation is also increased at higher cell densities, so cells were typically seeded at ~80-90% density on day 0, with attention that all cell lines in the experiment were seeded at the same density. Cells in low serum proliferate very slowly, if at all, so densities approached confluence without overcrowding that would challenge scoring.

### CRISPR-Cas9 genome editing

Genome editing in MEFs was performed as previously described (25,34,35). Benchling software (www.benchling.com/academic/) was used to design four 20 nt guides. To facilitate expression from the U6 promoter, a “G” was substituted for the first nucleotide for each guide RNA. Primers were purchased from IDT based on the following templates: 5′-CACC(N_20_)-3′ and 5′-AAAC(NR_20_)-3′, where N_20_ and NR20 refer to the 20 nt protospacer sequence and its reverse complement, respectively. The guides used to generate KO lines in MEFs were as follows.

Elmod1 guide 1: CACCGGATGCGGAAACTCACCGGA

Elmod1 guide 2: CACCGTTTGCTACGGCACCAAACC

Elmod3 guide 2: CACCGATGCCATGGTTCGTCAGCT

Elmod3 guide 3: CACCGCCCATTGGTTTCTGCCGTC

Complimentary oligos were annealed and cloned into pSpCas9(BB)-2A-Puro (PX459) V2.0 vector (Addgene plasmid #62988) at the *Bbs*I sites. Guides were targeted near the N-terminus of the protein and upstream of the ELMOD domain to optimize the likelihood of null alleles. We generated at least two different clones from at least two different guides, each with unique frameshifting mutations on both alleles, to protect against both off-target effects and the potential for use of downstream initiation of protein synthesis, alternative splicing, or other confounding changes (90). Two ELMOD1 guides were used to generate four independent ELMOD1 KO clones (two from each guide), while two ELMOD3 guides, targeting exons 5 and 8, were used to generate 13 clones (3 from one guide and 10 from the other) KO’d for ELMOD3. We also generated two ELMOD1/ELMOD3 double KO (DKO) lines by using one of the ELMOD3 guides (guide 2) transfected into one of our ELMOD1 KO lines (KO #2; Fig. S1C).

Low-passage MEFs were grown to 90% confluence in six-well dishes, transfected via Lipofectamine 2000 at a 1:3 ratio of DNA to Lipofectamine reagent with 4μg of DNA, and then replated onto 10 cm plates for growth overnight. Puromycin (3 μg/ml; Sigma #P8833) was added the next day and maintained for 4 days to enrich for transfected cells. Individual clones were isolated via limited dilution in 96-well plates, followed by expansion, cryopreservation, and sequencing of genomic DNA after PCR amplification of the region surrounding the targeted site to identify frameshifting mutations that propagate early stop codons. After initial testing of phenotypes and finding consistencies between KO lines, we chose two from each gene for more detailed analyses, though often examined every KO line available.

During the cloning of the *Elmod3* KO lines, but more so the DKO lines, we noted an apparent decrease in cell attachment to plates, evident as rounded cells after plating at low cell density. Prior treatment of plates with fibronectin alleviated this problem. To assess possible changes in cell attachment in the cloned lines, both the number and size of focal adhesions were quantified, and no differences were observed (Fig. S9A,B). Thus, an attachment defect was not evident in the cell lines described here, nor in cloning of lines KO’d for multiple other genes studied in our lab. Thus, the reasons for possible attachment problems are unknown and were not pursued further at this time.

### Flow cytometry analysis for DNA Content

As previously described (Turn et al., 2020), unsynchronized cells were prepared for flow cytometry by trypsinizing the cells, taking up the cells in ice cold PBS, washing with ice cold PBS, and fixing the cells with ice-cold 70% ethanol dropwise while simultaneously vortexing to reduce risk of cell clumping. Cells from both the supernatant and adherent cells were collected to ensure that we had a full representation of the cell population. The day of flow cytometry, cells were spun down, washed with ice-cold phosphate citrate buffer (0.1M citric acid in PBS, pH7.8) two times, treated with RNAse A for 15 minutes (100 μg/mL; Sigma; R5125), and treated with propidium iodide for 45 minutes (50 μg/mL; Sigma, P4170) to stain for DNA content. Cells were passed through a cell strainer and run on a FACSymphony A3. The G1 peak of WT cells was set at a 50K voltage for each run, and these settings were used to acquire all subsequent samples run that day to ensure that we accurately track 2N, 4N, and >4N peaks. At least 10,000 cells were collected per sample. FloJo software was used to plot data shown.

### Microscopy

Prior to imaging, cells were plated onto fibronectin coated 18 mm glass coverslips (#1.5; Fisher Scientific; 12-545-81), according to the specified experimental conditions, and processed for immunocytochemistry. Images were collected via widefield microscopy on our Olympus IX81 microscope with Slidebook software at 60X and 100X magnification (UPIanFI, 1.30 NA Oil). The same acquisition settings were used to ensure accurate comparisons (*e.g.*, same sample preparation, magnification, gain, offset, laser power, etc.). Data were processed using FIJI imaging software, making sure to apply the same image processing techniques to each sample of a given data set (*e.g.*, cropping, magnification, brightness, contrast, background subtraction, *etc.*). In the case of retinal tissue, samples were imaged using a Leica DM6000B deconvolution microscope (Leica Microsystems, Bensheim, Germany) or a Leica SP8 laser scanning confocal microscope (Leica Microsystems, Bensheim, Germany). Data were processed via Adobe Photoshop CS (Adobe Systems, San Jose, CA).

### Immunofluorescence

The following protocols were used based on the antigen we were targeting (described below):

*Methanol Fixation protocol*: (mostly for centrosomal/centriolar markers to provide cleaner imaging without cell background/preserve structure) Cells plated on coverslips were fixed with methanol for 10 minutes at −20°C before being washed 4x with PBS. Cells were blocked for 1hr at room temperature with 10% FBS in PBS and incubated overnight with primary antibody diluted in 10% FBS in PBS at 4°C. Cells were washed with PBS 4x before being incubated with secondary antibody (1:500 dilution in 10% FBS in PBS) for 1hr at room temperature in the dark. Cells were washed with PBS 2x, stained with 1:5000 Hoechst in PBS for 4 minutes, and washed 2x with PBS before being mounted overnight with a 1:9 ratio of PPD (p-phenylenediamine dihydrochloride; ACROS Organics; 624-18-0) to MOWIOL 4-88 Reagent (CALBIOCHEM; 475904) mounting medium.

*PFA Fixation protocol*: (to preserve membranes/ as a good general fixation protocol for cilia). Cells were fixed with prewarmed (37°C) in 4% PFA in PBS for 15 minutes. Cells were washed 2x with PBS and permeabilized with 0.1% Triton X-100 for 10 minutes. Cells were blocked with 1% BSA in PBS for 1hr at room temperature and incubated with primary antibody diluted in 1% BSA in PBS overnight at 4°C. Cells were washed 4x with PBS, incubated with secondary antibody diluted in 1% BSA in PBS for 1hr at room temperature, and washed and mounted on slides as described above.

Note that use of the same antibody but with different staining protocols may yield differences. For example, INPP5E at cilia is optimally visualized after PFA fixation, while its presence at Golgi is most prominent after methanol fixation.

### Immunoblotting

Cells were collected and cell pellets were lysed in PBS with 1% Triton X-100 on ice. After 15 min, cells were spun in a microfuge at 14,000xg for 30 min to remove insoluble material. Protein (40 μg) were loaded into lanes of an 11% polyacrylamide gel and resolved at 60 mA. Proteins were later transferred onto nitrocellulose filters overnight at 20 mV. The next day, filters were stained briefly with Ponceau S to confirm equal transfer, then put into Blotto blocking buffer (5% dry milk in PBS) for one hr before addition of primary antibody overnight. Filters were washed 3 × 10 min in PBST (PBS with 1% Tween-20) before adding secondary antibody, incubating for one hr, followed by 3 × 10 min washes in PBST. Imaging was performed using a Bio-Rad imager.

### Scoring of cell phenotypes

For all phenotypes described above, experiments were performed in triplicate and scored in at least duplicate, 100 cells per a sample. For phenotypes such as ciliation, nucleation, and centrosome/centriole counts, scoring was binned based on the number of each organelle present/ the number of the organelles positive for that marker. For markers in which it was related to a degree of localization in cilia (e.g., ARL13B), we binned them as either present (visible even without checking the Ac Tub channel), reduced (present, but only noticeable upon switching to Ac Tub channel), or absent (cannot be detected even upon switching to Ac Tub channel). For all ciliary phenotyping, Ac Tub was used as a co-stain to detect cilia and draw accurate comparisons. For centrosomal/basal body scoring, γ-tubulin was used as the standard comparison point. Finally, for Golgi staining/localization, either GM130 or Golgin97 was used to mark Golgi.

### Sample preparation and staining of mouse retinae

*Animals:* Transgenic eGFP-CETN2 mice (155) were kept on a 12-hour light-dark schedule at 22°C, with free access to food and water. Animal health was monitored on a regular basis, and all other procedures complied with the German Law on Animal Protection and the Institute for Laboratory Animal Research Guide for the Care and Use of Laboratory Animals.

*Immunohistochemistry of retinal sections*. Mouse retinae were dissected from enucleated eye balls and cryofixed in melting isopentane and cryosectioned as previously described (156,157). Cryosections (10 μm thick) were placed on poly-L-lysine-precoated coverslips and incubated with 0.01% Tween 20 in PBS for 20 min. After washing, sections were flooded with blocking solution (0.5% cold-water fish gelatin plus 0.1% ovalbumin in PBS) and incubated for at least 30 min followed by an overnight incubation with primary antibodies at 4°C in blocking solution (91). Washed cryosections were then incubated with secondary antibodies conjugated to Alexa 488 or Alexa 568 (Invitrogen) in blocking solution and with DAPI (Sigma-Aldrich) to stain the DNA of nuclei, for 1.5 h at room temperature in the dark. After three washes in PBS, specimens were mounted in MOWIOL 4.88 (Hoechst) and imaged using a Leica DM6000B microscope (Leica).

### Statistics

All experiments were scored at least in duplicate and performed at least in triplicate, using at least two clones from at least two different guides for each gene targeted (ELMOD1 or ELMOD3). Unless otherwise stated, at least 100 cells were scored per sample. Error bars in the graphs represent the standard error of the mean (SEM), and box-and-whisker plots indicate the range of the data along with the median and upper/lower quartiles. One-way or two-way analysis of variance (ANOVA) tests were used to determine whether there were significant differences between test groups. The presence of an asterisk in a figure indicates statistical significance: \**p* < 0.05, as also indicated in figure legends. We consider the individual lines as biological replicates. Therefore, if we report that a sample has an *N* = 4, this indicates that four different lines were scored in (at least) duplicate, and the averages of those duplicates are presented in the graphs.

## ACKNOWLEDGMENTS

This work was supported by grants from the National Institutes of Health: R35GM122568 to R.A.K., R35GM122549 to T.C., 1F31CA236493-02 to R.E.T, 1F31HD096815-03 to S.I.D, the Foundation Fighting Blindness (FFB PPA-0717-0719-RAD) to U.W., the German Research Council/DFG in the framework of the SPP SPP2127 - Gene and Cell based therapies to counteract neuro-retinal degeneration WO548/9-1 to U.W., R01GM127361 to J.E.C., and the joint training program between Emory University School of Medicine and Xiangya School of Medicine, Central South University (Y.H.). We thank colleagues for their generous sharing of key reagents, in particular Rytis Prekeris and Jim Goldenring for antibodies and plasmids. This research was supported in part by the Emory University Integrated Cellular Imaging (ICI) Microscopy and Pediatrics Flow Cytometry Cores. We thank the Emory Biochemistry Department and Laney Graduate School for their ongoing support and shared resources provided.

---

**Figure S1:**
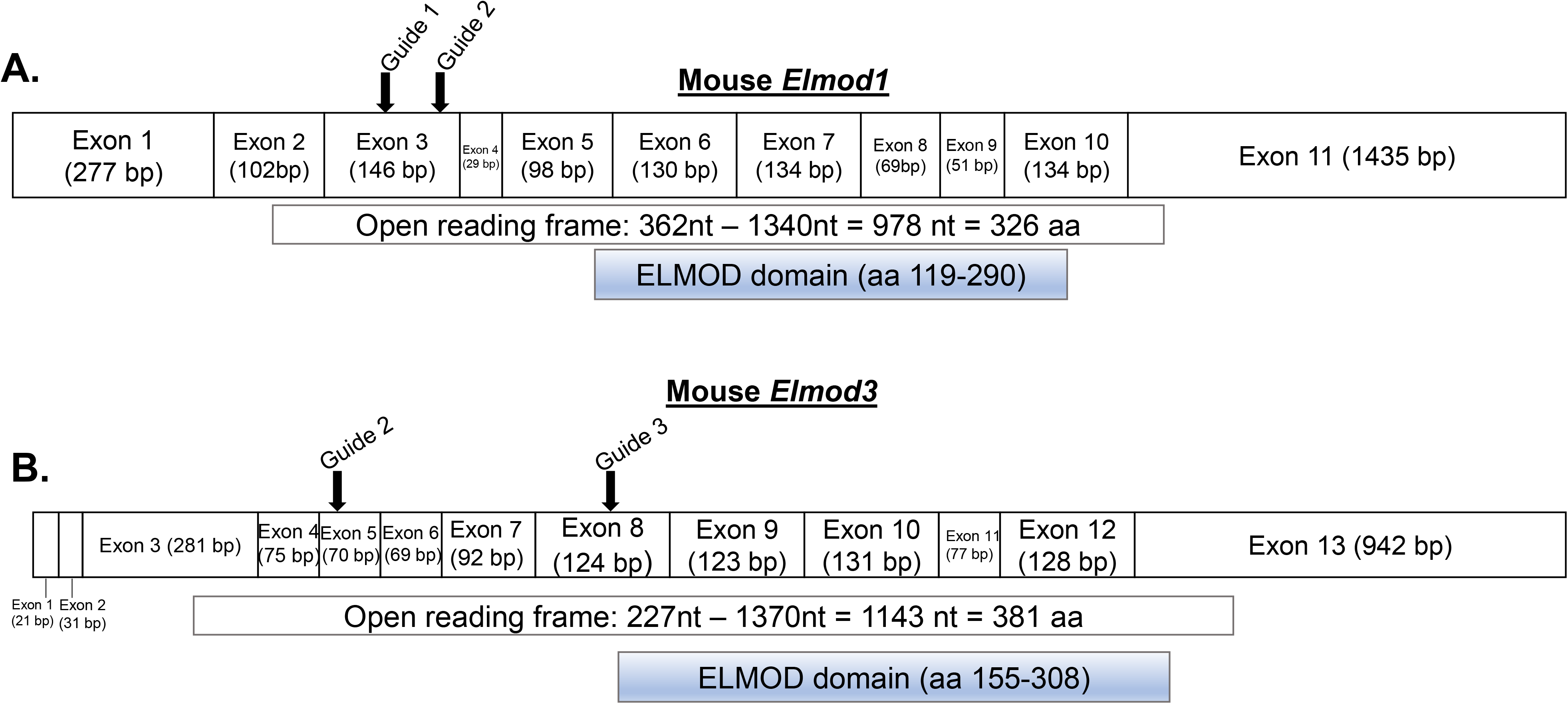
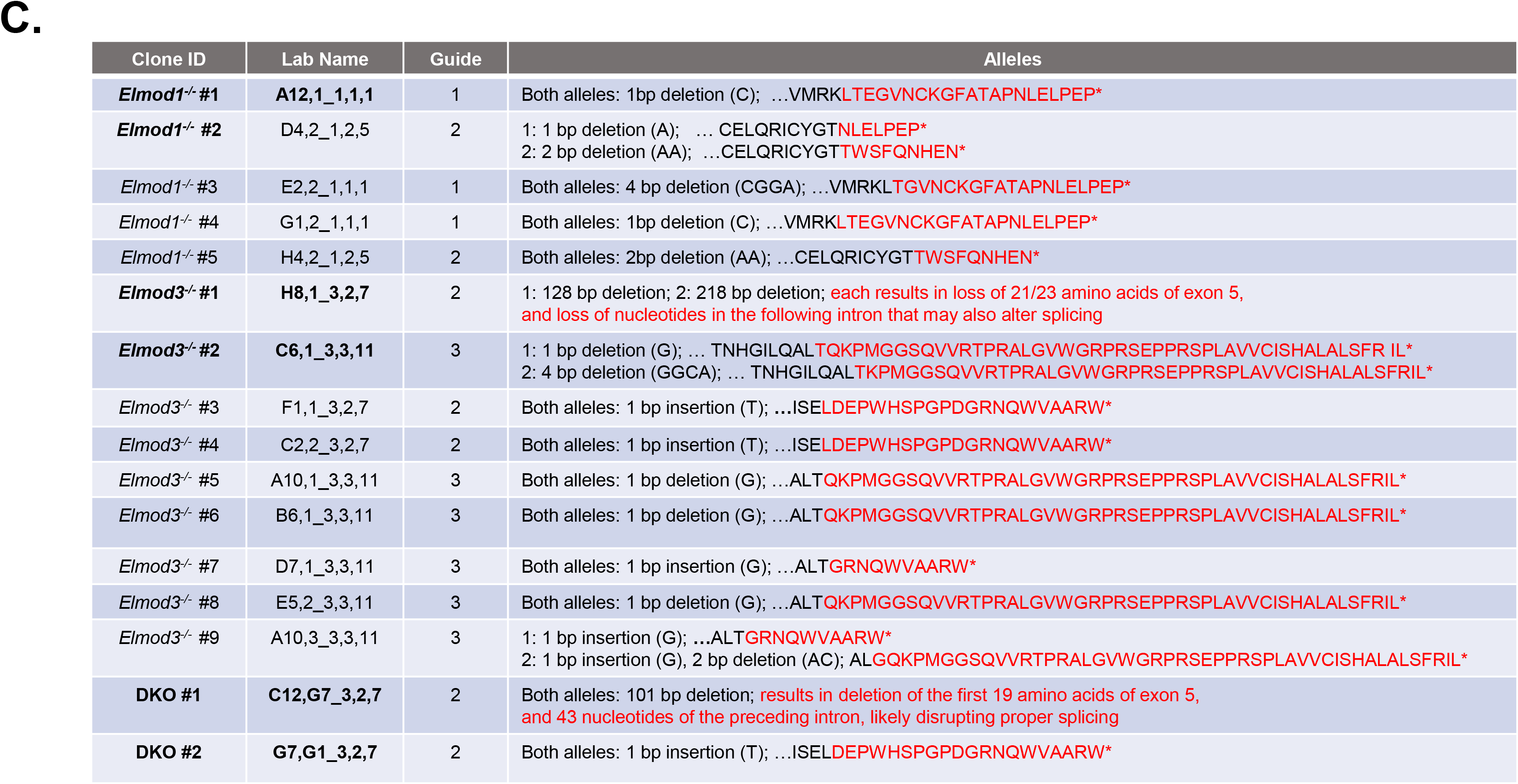
Generation of *Elmod1* KO, *Elmod3* KO, and DKO MEFs. (**A, B**) Diagrams show the open reading frames encoding ELMOD1 and ELMOD3 proteins under the exons and where the guides target in the exons and corresponding protein sequences. Note that all the guides generated were made to target upstream of the ELMOD domain, increasing the likelihood that if any protein product is made, it is non-functional. The locations of the ELMOD domains are shown under the open reading frames. (**C**) The names and alleles of each of the cloned KO lines used are shown. The two *Elmod1* KO, two *Elmod3* KO, and two DKO lines used in all studies are indicated in bold, though all cloned KO lines are shown here as all were tested in at least some of the reported assays and yielded the same results as the two studied most thoroughly.

**Figure S2:**
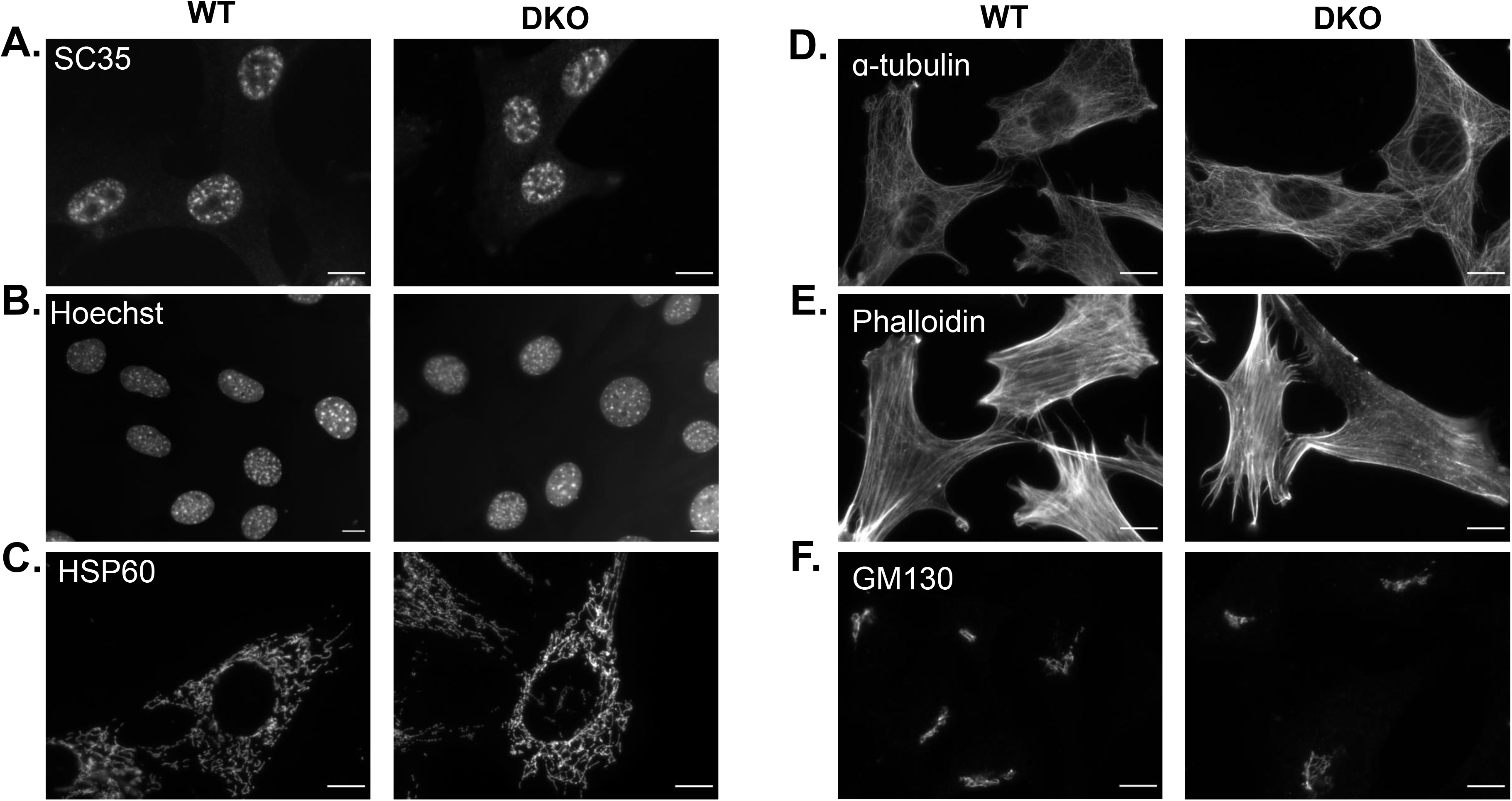
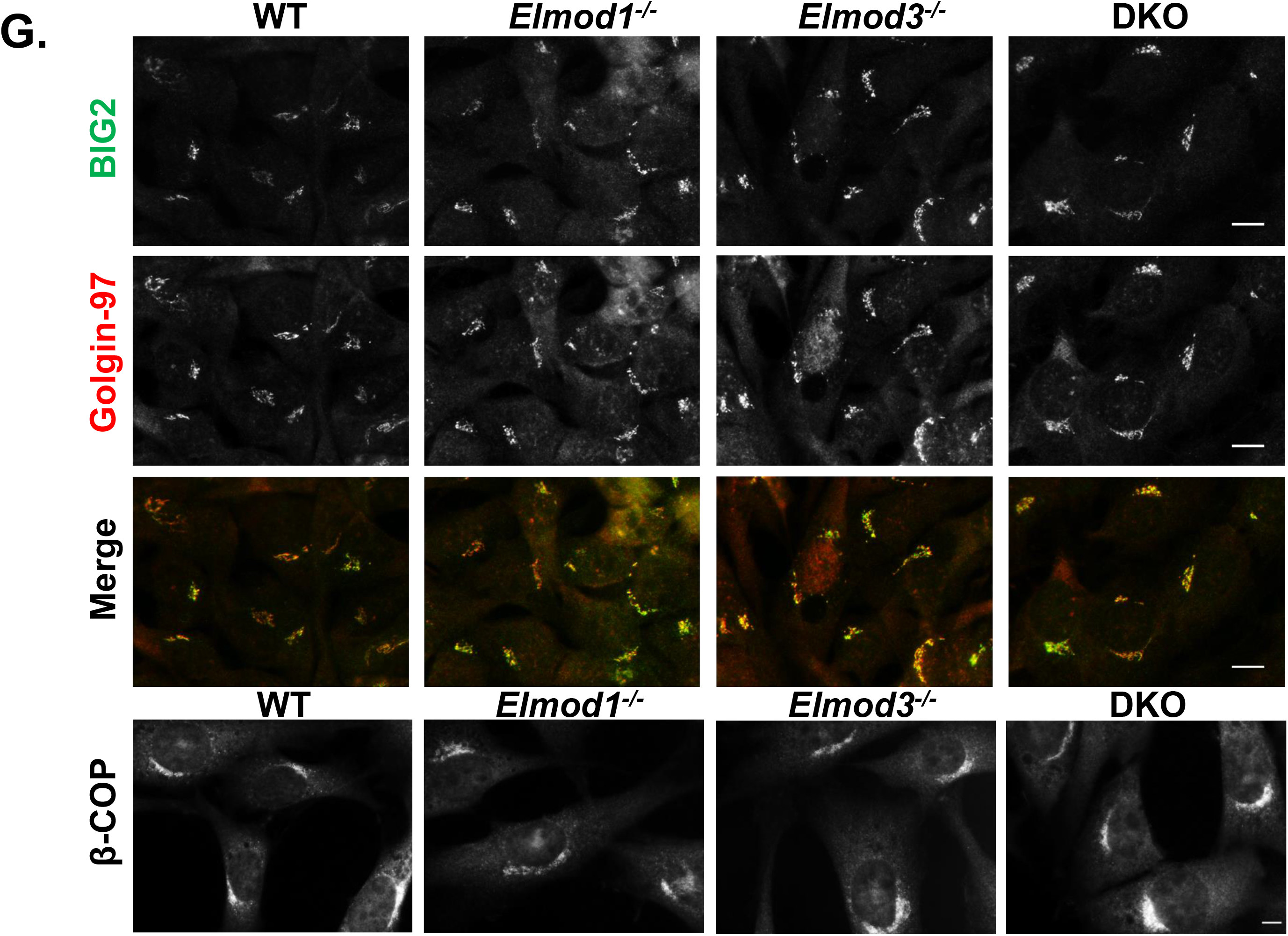
*Elmod1*, *Elmod3*, or DKO MEFs display no evidence of changes in staining of nuclear speckles (A: SC35), nuclei (B: Hoechst), mitochondria (C: HSP60), microtubules (D: α-tubulin), f-actin (E: phalloidin), or Golgi (F: GM130). (**G**) Because of later evidence of changes in traffic of specific proteins from the Golgi, we examined other Golgi markers and again observed no differences between WT and KO cells. Here, cells were co-stained for the ARF GEF BIG2 (aka ARFGEF2) and Golgin-97, or separately for β-COP (bottom panels). Cells were fixed with 4% PFA, permeabilized with 0.1% Triton X-100, and stained for the indicated markers in two lines each of WT, *Elmod1* KO, *Elmod3* KO, and DKO. Screening was repeated at least in triplicate. Representative images of WT and KO cells were collected at 100X magnification using widefield microscopy, and images were processed via FIJI imaging software. Scale bar = 10 μm.

**Figure S3:**
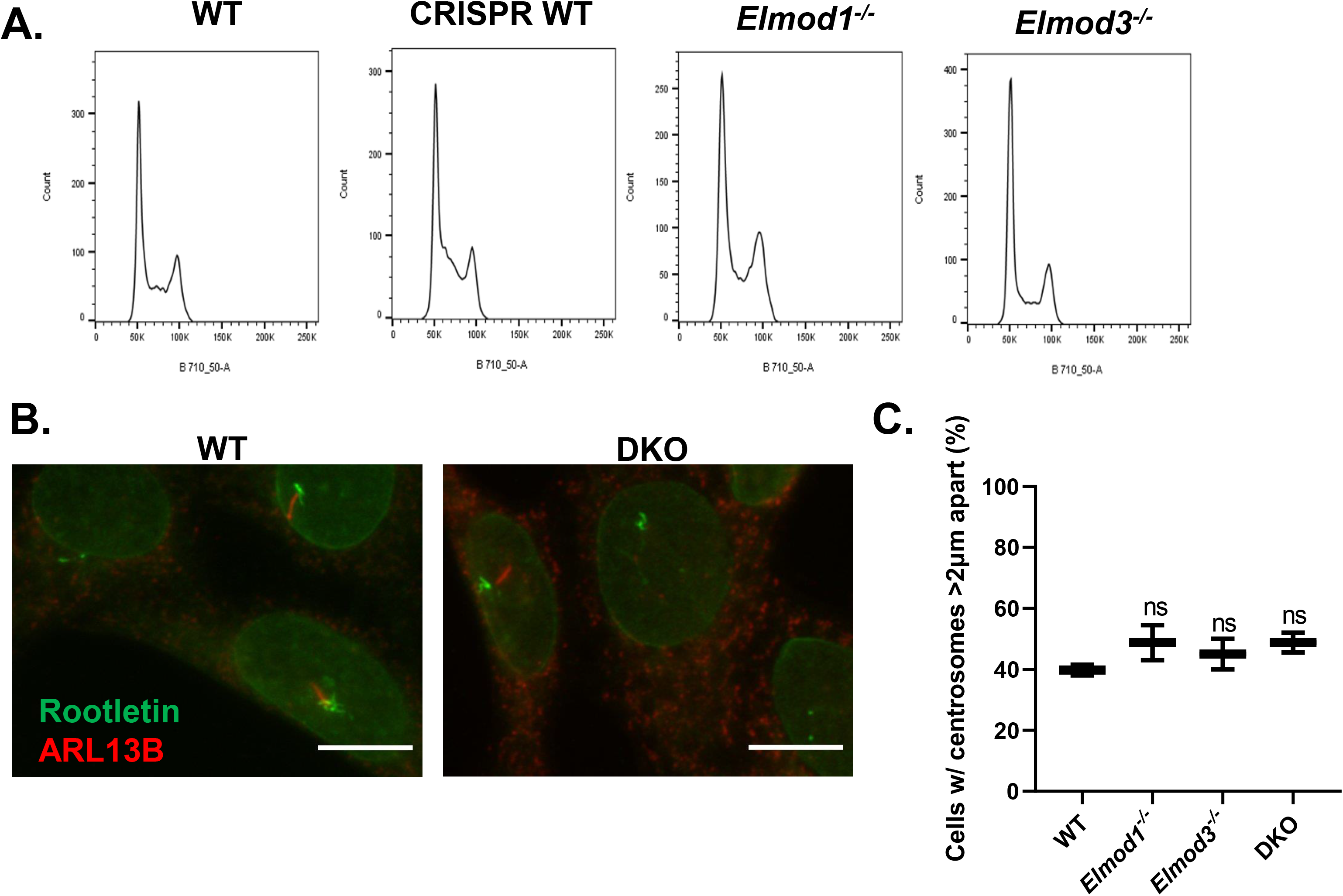
*Elmod1 KO*, *Elmod3 KO*, or DKO MEFs exhibit no evident defects in cell cycle, rootlet morphology, or centrosome separation. **(A)** Two lines each of WT, *Elmod1* KO, *Elmod3* KO, and DKO MEFs growing in log phase were collected, stained for propidium iodide, and examined for DNA content by flow cytometry, as described under Methods. At least 10,000 cells were analyzed per sample, and data were processed via FloJo software. This experiment was repeated in triplicate with each of the eight cell lines, and no consistent differences were evident. Representative graphs are shown. **(B)** The same eight cell lines described in **(A)** were serum starved for 24 hrs prior to fixation and were co-stained for rootletin and ARL13B. The experiment was repeated at least three times for each cell line. Images of WT and DKO cells are shown were collected at 100X magnification using widefield microscopy. Note that one of the DKO cells shown displays ARL13B staining in its cilium, to highlight that while ciliation and ARL13B staining of cilia are each strongly reduced, each can be found. Scale bar = 10 μm. **(C)** These same cell lines were grown to ~75% confluence, fixed, and stained for γ-tubulin to mark centrosomes. Images were collected, and FIJI measuring tool was used to quantify the percentage of cells with centrosomes farther than 2μm apart. Results were plotted in GraphPad software, with error bars indicating SEM. ns = p>0.05, calculated via One-Way ANOVA.

**Figure S4:**
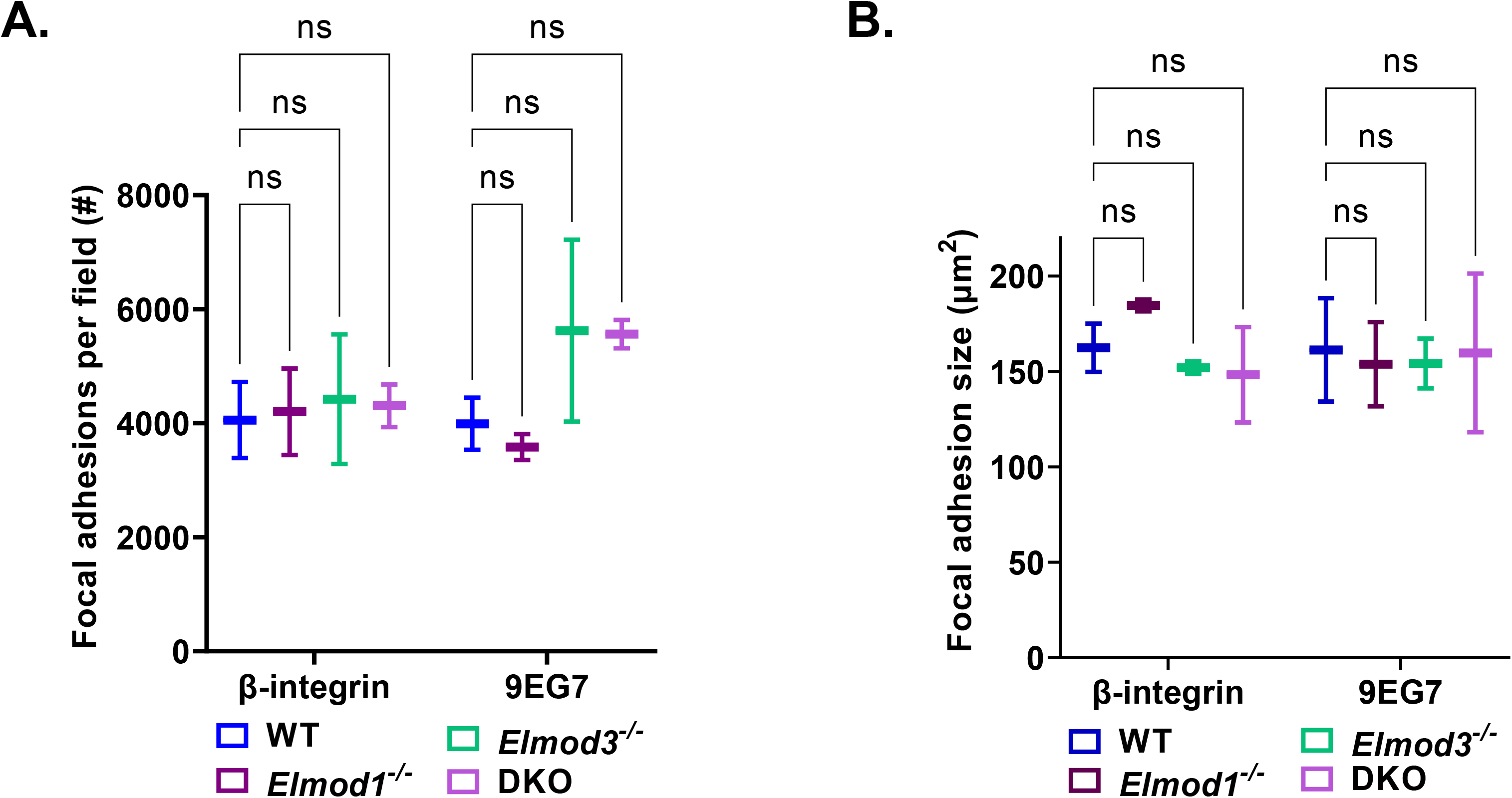
ELMOD1-myc is expressed to much higher levels than is ELMOD1-myc in MEFs. WT cells were transfected with plasmids directing the expression of either empty vector, ELMOD1-myc, or ELMOD3-myc, as indicated above each lane. The next day, cells were collected, and total cell lysates were generated, as described under Methods. Equal protein (40 μg/lane) was loaded on each lane of a 13% polyacrylamide gel and later transferred onto nitrocellulose membranes for 90 min at 60V. The membrane was then stained with Ponceau S (left panel) or with anti-myc antibody and developed using enhanced chemiluminescence (right panel). Molecular weight standards were run in a separate lane, and sizes are indicated between panels.

**Figure S5:**
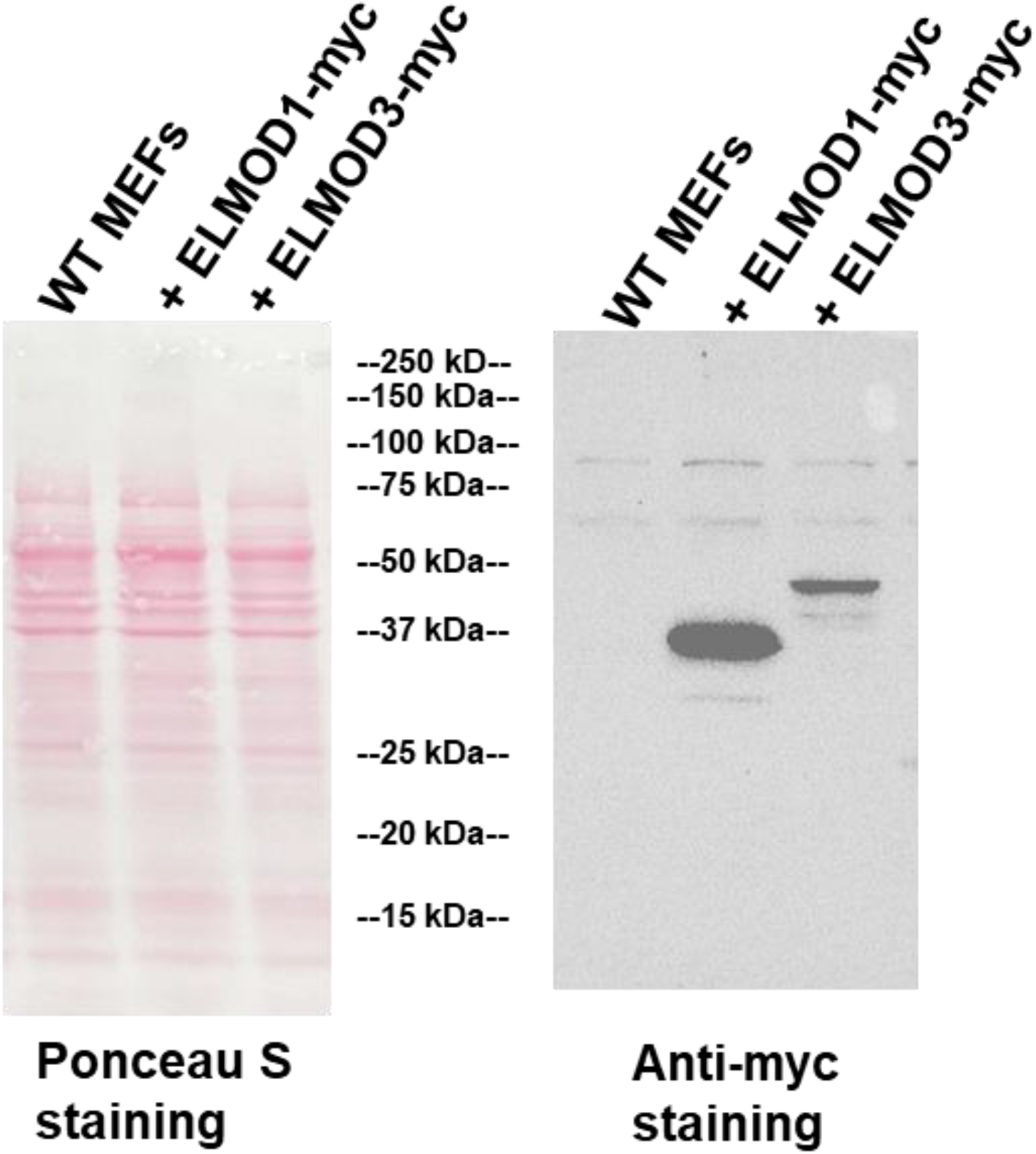
Neither *Elmod1* nor *Elmod3* KO alters the extent of CEP164 recruitment and CP110 release from the basal body, despite the decrease in ciliation. Two lines each of WT, *Elmod1* KO, *Elmod3* KO, and DKO cells were serum starved for 24 hours before fixation and staining for either (**A**) CEP164 or (**B**) CP110 and γ-tubulin, as described under Methods. The presence of CEP164 and CP110 were scored in 100 cells per cell line in duplicate, binning cells as having either 0, 1, or 2 centrosomes positive for CEP164 or CP110. Data were graphed using GraphPad Prism. Error bars = SEM. (**C**) Representative widefield images of cilia from cells stained for CEP290 and Ac Tub are shown, with the genotypes of the cells shown at the top. Scale bar = 10 μm.

**Figure S6:**
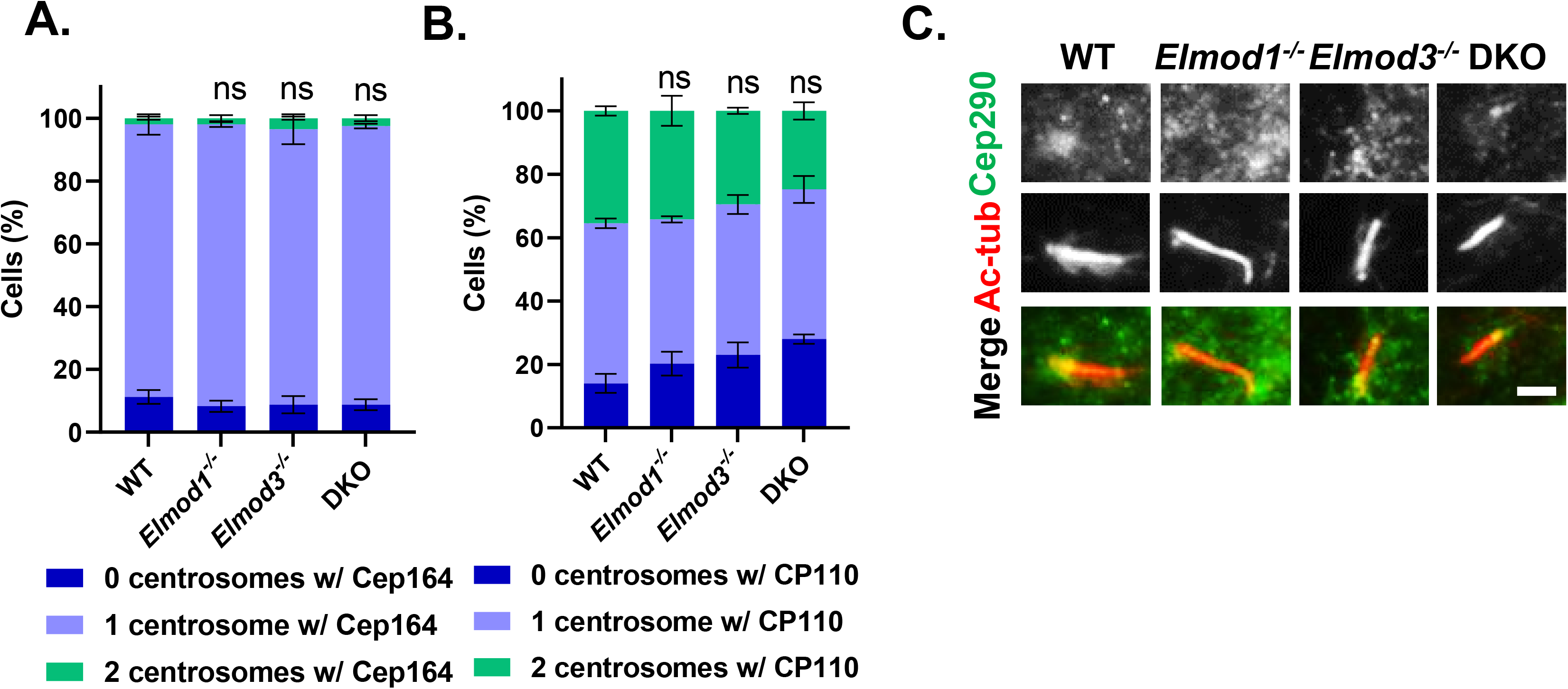
Gli3 recruitment to cilia is unaffected by deletion of *Elmod1* or *Elmod3*. The standard eight cell lines were serum starved for 24 hr before fixation and co-staining for Gli3 and Ac Tub. This experiment was repeated at least in triplicate with consistent lack of changes. Representative images from each genotype were collected at 100x magnification via widefield microscopy. Scale bar = 10 μm.

**Figure S7:**
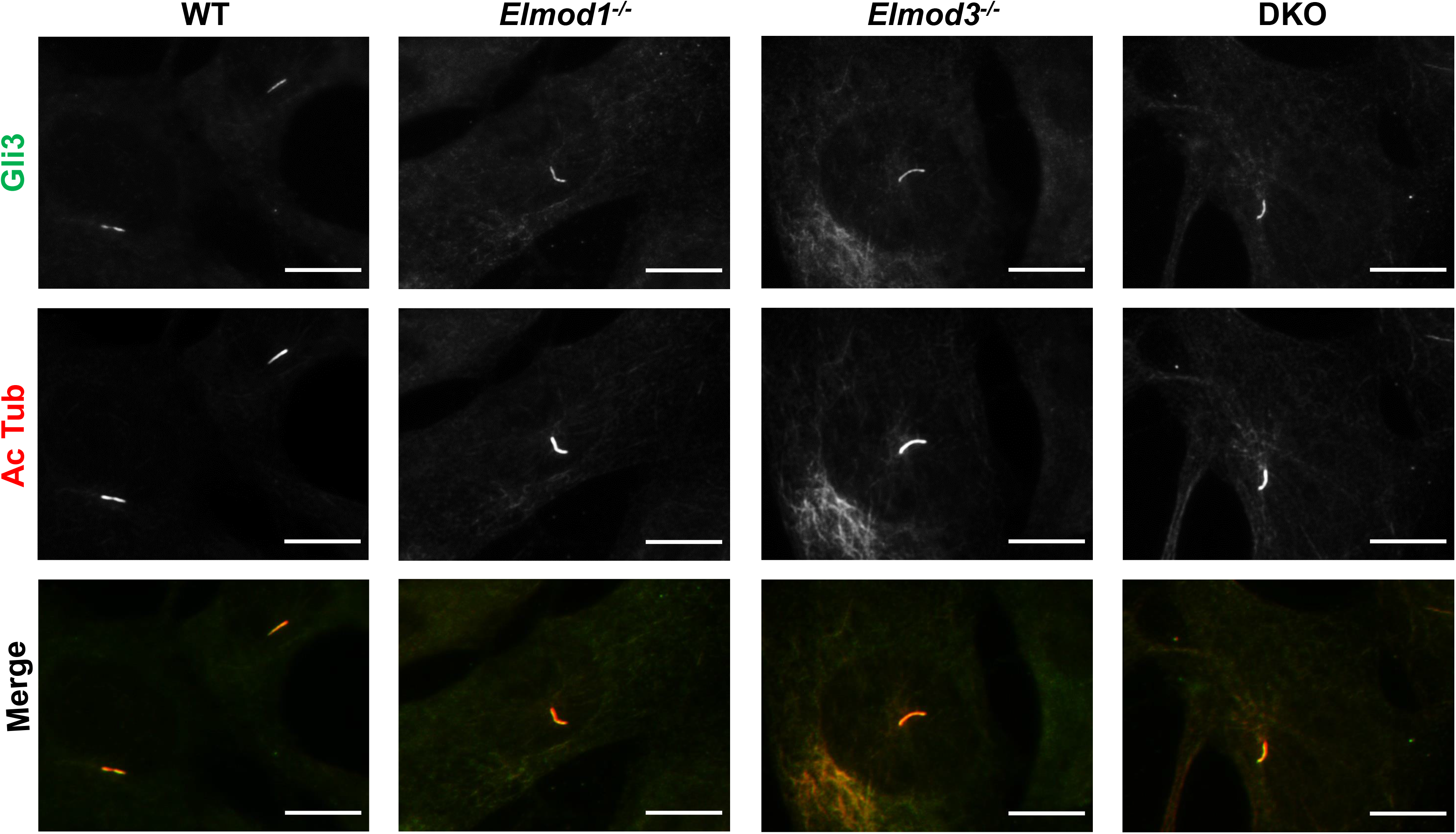
IFT140 colocalizes with Rootletin in both WT and KO cells. The standard eight cell lines were serum starved for 24 hours before being fixed with ice-cold methanol and stained for IFT140 and rootletin (to mark the ciliary rootlet). Representative images from each genotype were collected at 100x magnification via widefield microscopy. Scale bar = 10 μm.

**Figure S8:**
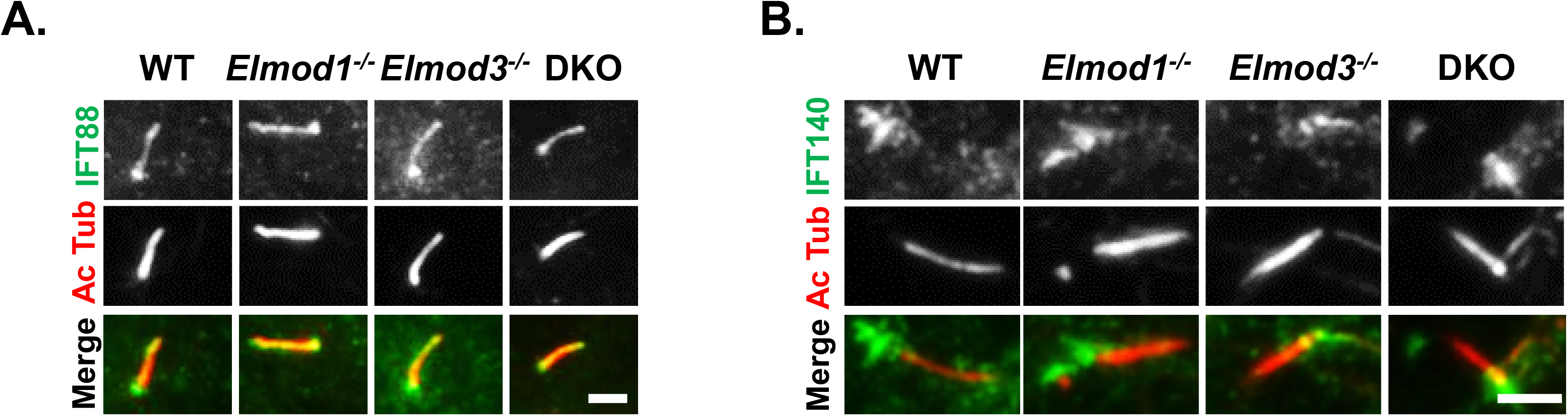
IFT88 and IFT140 localization to cilia in ELMOD1/3 KO lines is unchanged. Eight working cell lines that were serum-starved for 24 hours were fixed with 4% PFA and stained for Ac Tub (to mark cilia) and either IFT88 (**A**) or IFT140 (**B**). Representative images from each genotype were collected at 100x magnification via widefield microscopy. Scale bar = 10 μm.

**Figure S9:**
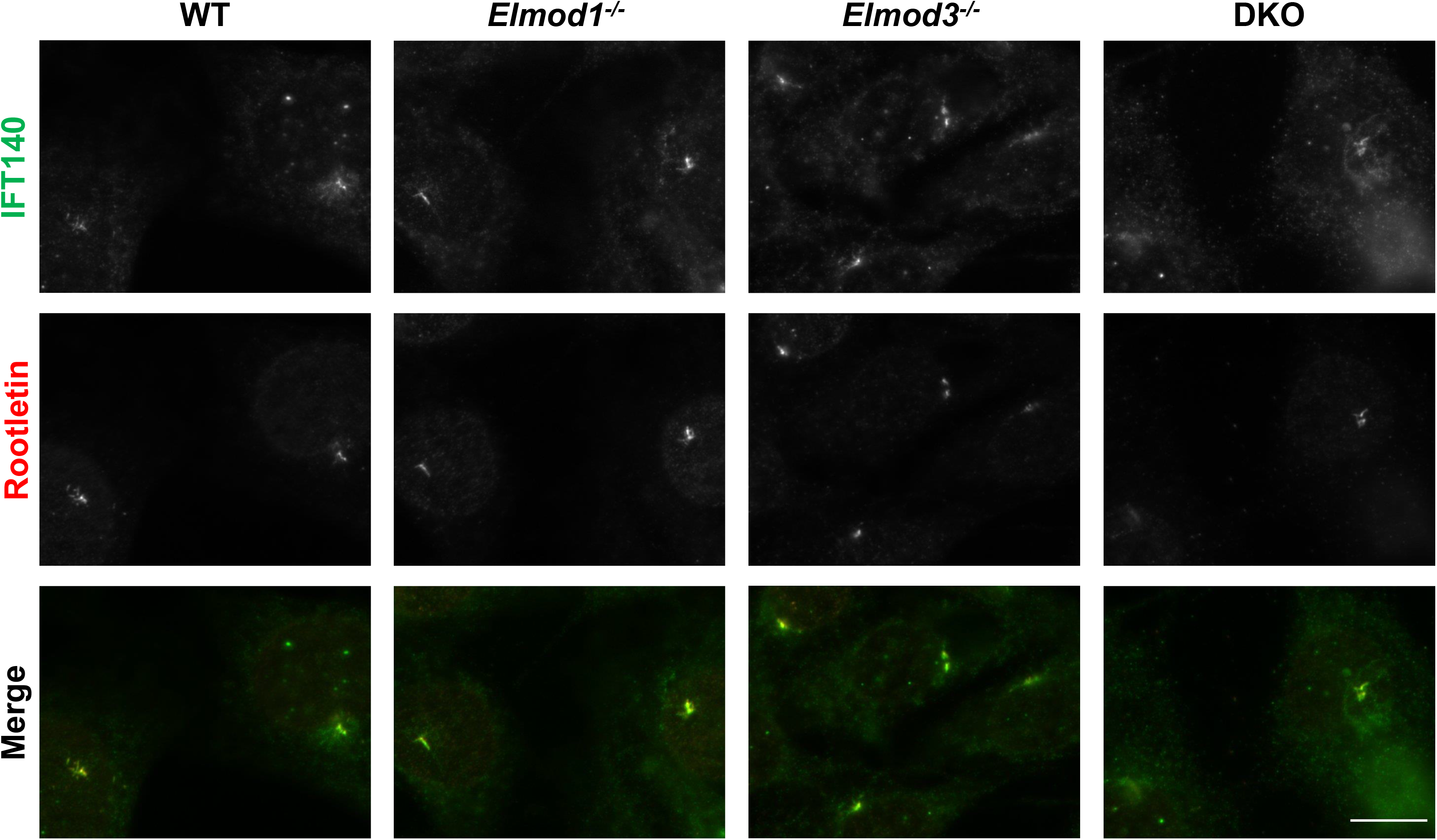
Number and sizes of focal adhesions are unchanged in ELMOD1 and ELMOD3 KO lines. The eight working cell lines were tested for changes in focal adhesion architecture via immunofluorescence, staining for total β-integrin and active (ligand-bound) β-integrin using the activation-specific antibody 9EG7. Image quantification was performed using FIJI software, applying the same conditions to subtract background and to set thresholds for each sample. Data were processed via GraphPad prism software, and statistical significance was assessed via One-Way ANOVA. Error bars = SEM. * = p<0.05 as calculated by One-Way ANOVA.

## Footnotes

1 Abbreviations used in the text include: Ac Tub, acetylated tubulin; CWT, CRISPR-WT (CRISPR-treated cells without edits in the targeted region); DKO, double knock out (of both ELMOD1 and ELMOD3); FBS, fetal bovine serum; GAP, GTPase activating protein; GEF, guanine nucleotide exchange factor; GFP, green fluorescent protein; IFT, intraflagellar transport; KO, knockout; MEF, mouse embryonic fibroblast; PCM, pericentriolar material; PFA, paraformaldehyde; SHH, Sonic Hedgehog; Smo, Smoothened; TZ, transition zone; WT, wild-type.

